# Neuronal avalanches as a predictive biomarker for guiding tailored BCI training programs

**DOI:** 10.1101/2025.05.31.657206

**Authors:** Camilla Mannino, Pierpaolo Sorrentino, Mario Chavez, Marie-Constance Corsi

## Abstract

Motor imagery-based Brain-Computer Interfaces (BCIs) restore control in persons with motor impairments, but up to 30% of users struggle—a phenomenon known as “BCI inefficiency”. This study tackles a key limitation of current protocol: the use of fixed-length sessions training paradigms that ignore individual learning variability. We propose a novel approach based on *neuronal avalanches*— spatiotemporal cascades of brain activities—as biomarkers to characterize and predict user-specific learning. From electroencephalography data across four sessions in 20 subjects, we characterized avalanches by their length and their spatiotemporal size. These features showed significant training and task effects and were found to correlate to BCI performance across sessions. We further assessed their ability to predict BCI success through longitudinal models, achieving up to 91% accuracy, improved by spatial filtering on selected brain regions. These findings demonstrate the utility of neuronal avalanche dynamics as robust biomarkers for BCI training, supporting the development of personalized protocols aimed at mitigating BCI illiteracy.

## Introduction

Brain-Computer Interfaces (BCIs) are promising technologies for restoring communication and control in individuals with severe motor and speech impairments. Despite their potential, using a non-invasive technique, a persistent limitation remains: a substantial proportion of users—estimated at 15–30%—fails to gain effective control over the interface, a phenomenon commonly referred to as “BCI inefficiency”^1^. One underappreciated contributor to this challenge lies in the uniform design of BCI training protocols, which typically prescribe a fixed number of sessions for all participants. This “one-size-fits-all” approach overlooks the individual mechanisms underlying BCI performance and neural learning, assuming a homogeneous neural learning across users. Considerable inter-individual variability exists in neural plasticity^2^, cognitive strategy^3^, and brain dynamics^4^, which implies that users follow distinct learning trajectories^5^. While some individuals quickly internalize the required mental strategies, others may require longer or more personalized training to induce the necessary neurophysiological adaptations^6^.

A growing body of evidence suggests that both psychological and neurophysiological factors contribute to BCI performance variation, manifesting both across users (inter-subject variability)^7^ and within individuals over time (intra-subject variability)^7^. Human factors and cognitive traits such as motivation, emotional state, or the strategy use modulate BCI performance.^8^ As an example, higher motivation and positive mood have been associated with enhanced performance^9,10^, while frustration and overconfidence can impair^11,12^. Passive or emotionally-driven strategies tend to outperform effortful cognitive techniques^4,3,5^, and traits such as visuo-motor coordination, attentional control, imagination ability, and musical training have been linked to better outcomes ^13,14^.

On the neurophysiological side, several biomarkers based on spatially localized features, like event-related desynchronization/synchronization (ERD/ERS), event-related potentials (ERP) and sensorimotor rhythms (SMR), have been proposed to predict BCI aptitude. Resting-state electroencephalography (EEG) markers, notably higher alpha power, are consistently associated with better MI control, whereas BCI-illiterate individuals often show increased theta and decreased alpha activities^15,16^. Additionally, fluctuations in gamma and alpha band activity have been linked to moment-to-moment variations in attentional engagement, further influencing task success ^17^. ERD, a core signature of motor imagery, tends to be diminished in low-performing users^18^, while dynamic features such as fluctuations in frontal gamma and preparatory alpha power reflect intra-individual variability across sessions^19,20^. Importantly, early training performance may itself serve as a predictive marker: Neumann and Birbaumer (2003)^21^, and Kübler at al., 2004^22^, showed that early slow cortical potentials (SCP) modulation correlated with later performance in healthy participants and severely paralyzed patients. Similarly, Halder et al, demonstrated that an auditory oddball response recorded before the P300 BCI session can be used to predict performance^23,24^. Weber et al. 2011^25^ found that SMR modulation success in early sessions predicted long-term outcomes, although reliable prediction required at least eleven sessions.

While these findings highlight the utility of preparatory activity and resting-state EEG features, a major limitation of current BCI paradigms is their reliance on univariate, localized measures—treating brain regions as isolated signal sources. This perspective neglects the fact that brain function emerges from dynamic interactions across distributed neural networks. Accordingly, functional connectivity (FC), defined as the statistical interdependence between brain signals generated from different regions, has gained attention as a system-level predictor of BCI performance^26,27^. For instance, Corsi et al., 2020^28^ reported that high-performing users progressively show stronger sensorimotor activation and decreased connectivity in associative cortical areas—indicating a transition toward more automatic and efficient BCI control. Similarly, Stiso et al., 2020^29^ demonstrated that task-related changes in functional connectivity, particularly within frontal networks, were associated with improved attentional modulation and performance. Complementary evidence from fMRI studies supports these observations, indicating that functional activation in regions such as the supplementary motor area and the parietal cortex distinguishes high from low performers^30,31^. Moreover, structural features, including white matter connectivity and the volume of the mid-cingulate cortex, have been found to correlate with BCI performance^32^. However, challenges remain in capturing the temporally evolving nature of these interregional interactions^26^.

Most BCI studies have historically focused on periodic, oscillatory signals—such as mu and beta rhythms—generated by synchronous neural activity. In contrast, aperiodic signals have often been dismissed as background noise, despite evidence suggesting that they encode essential information about the underlying state of neural populations^33^. Recent research indicates that both sustained oscillations and transient burst events coexist in the brain, each providing unique insights into cognitive and motor processes^34^. In particular, aperiodic bursts are starting to be understood as intermitten^35^. A compelling framework for capturing these dynamics involves neuronal avalanches—cascades of burst activity that propagate across cortical networks^36,37^. Shriki et al., 2013^38^ demonstrated that they might be observable at the scale of the entire human cortex with non-invasive neuroimaging methods. Indeed, using magnetoencephalography (MEG), they showed that resting state activity of healthy human subjects is organized in terms of neuronal avalanches. Detected by identifying large signal excursions in coarse-sampled EEG or MEG data, neuronal avalanches offer a spatiotemporal representation of how neural activity initiates and dissipates across brain regions^39^. These cascades have been shown to preferentially travel along white matter tracts ^40^, with a velocity dependent upon structural features of each bundle, and to shape spontaneous fluctuations in resting-state brain activity^41^. Recent work has demonstrated that features derived from the Avalanche Transition Matrices (ATMs)—which encode the probability of activity propagation between brain regions—can reliably distinguish between motor imagery and rest conditions in BCI tasks. These features have shown improved inter-subject consistency and better interpretability than conventional oscillatory power metrics^42,43^. Unlike EEG traditional features, which reflect the magnitude of localized activity, avalanche-based metrics capture how activity propagates across neural circuits over time—providing a network-level, dynamical account of brain function. This distinction is crucial for understanding individual variability in learning and performance. Moreover, recently the slope of the power spectral density, a parameter of the non-periodic, arrhythmic, and scale-free activity of the brain, has been proposed as predictor of visual p300-BCI from resting-state ^44^.

Building on the idea that learning induces measurable changes in brain activity, we investigated whether the dynamics of neuronal avalanches can serve as robust biomarkers to capture these changes and guide the development of personalized training programs in MI-based BCI training across sessions. We hypothesized that features derived from avalanche propagation— namely, avalanche length and spatial activation—encode both task-specific effects and longitudinal changes associated with learning. By capturing the evolution of brain-wide activity patterns, these features may reveal whether a user is adapting effectively and could therefore support early prediction of future performance. Ultimately, this approach seeks to move beyond standardized protocols and toward neurophysiologically informed, individualized BCI systems designed to guide personalized BCI training programs and overcome BCI illiteracy.

## Materials and Methods

### 2.1 Participants, Experimental Protocol, and EEG Acquisition

Twenty healthy, right-handed adults (mean age: 27.5 ± 4.0 years; 12 males), all BCI-naive and free of any neurological or psychological disorders, participated in the study. According to the declaration of Helsinki, written informed consent was obtained from subjects after explanation of the study, which was approved by the ethical committee CPP-IDF-VI of Paris. All participants received financial compensation for their participation. The experiment followed a longitudinal design, consisting of four EEG-based BCI training sessions over two weeks (two sessions per week). Between sessions, participants were instructed to continue training independently at home using short instructional videos.

The BCI task used in this study was a one-dimensional, two-target cursor control paradigm. To move the cursor toward the upper target, participants performed sustained motor imagery (MI) of right-hand grasping; to reach the lower target, they remained at rest, randomly and equally distributed between upper and lower positions. Each trial began with a 1-second inter-stimulus interval (ISI), followed by a 5-second target presentation. Visual feedback was displayed between 3 and 6 seconds, showing a cursor that started in the centre-left of the screen and moved rightward at constant speed. Participants were tasked with controlling the vertical position of the cursor through brain activity modulation *(Figure 1a)*.

**Figure 1:**
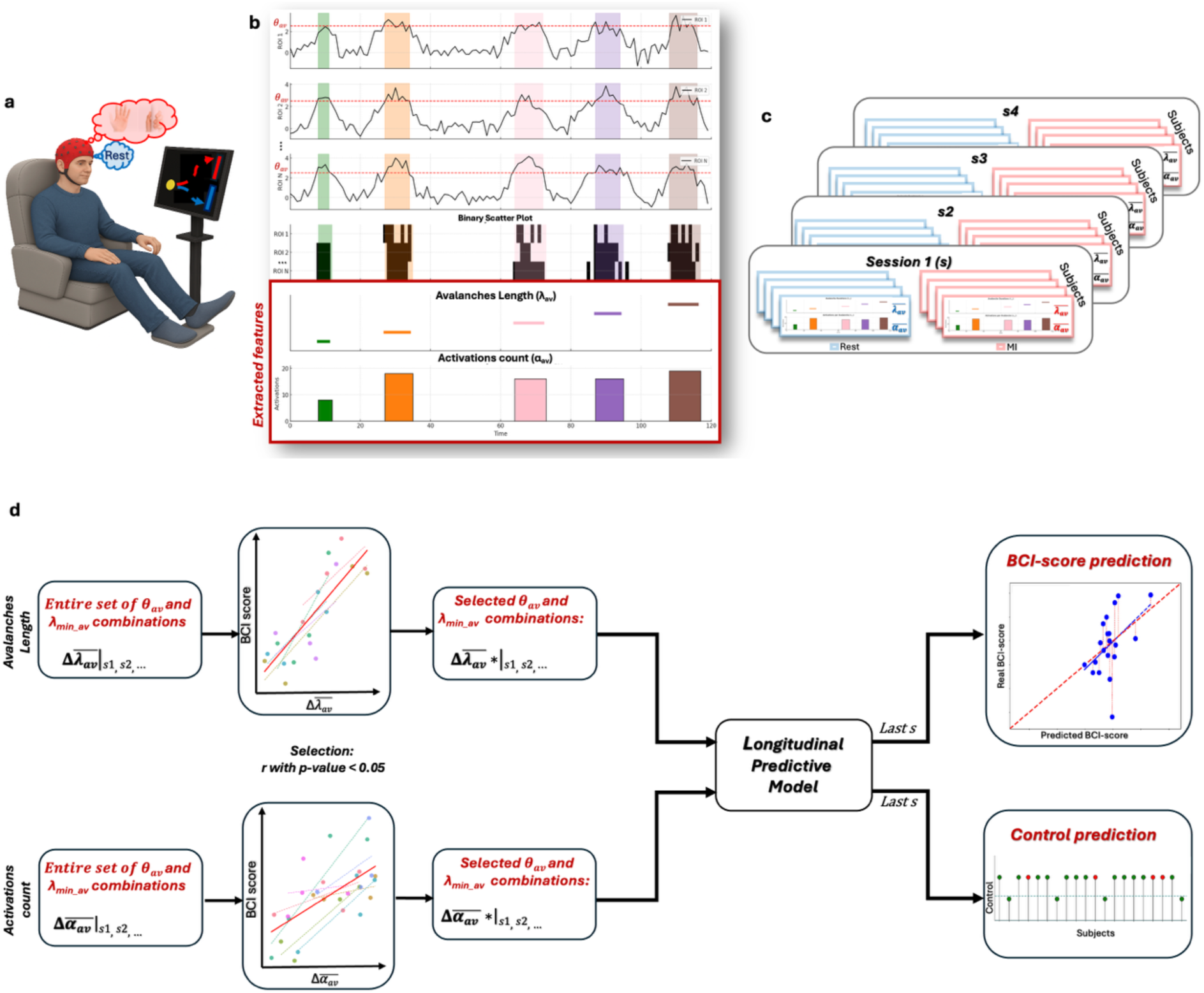
General analysis. **a)** Schematic representation of the experimental BCI protocol with EEG acquisition. **b)** Neuronal avalanche definitions across different ROIs are shown in the first three rows. The binary scatter plot derived from these definitions appears in the subsequent row. Feature extraction for λ_av_ and α_av_ is illustrated in the penultimate and last rows, respectively. **c)** Repetition of the analysis across different sessions (Session 1 to Session 4) and under two experimental conditions (Resting vs. Motor Imagery). **d)** Analysis workflow from extracted features (left blocks) to longitudinal predictive models (right blocks), including a regressive model (top right) and a classification model (bottom right). The transition through central blocks reflects parameter pair selection based on significance from repeated correlation analyses.

EEG signals were recorded using a 74-channel Easycap system (Ag/AgCl electrodes) arranged according to the international 10–10 system. Data acquisition took place in a magnetically shielded room with a sampling rate of 1 kHz and a 0.01–300 Hz bandpass filter. Artifact correction was performed using ICA (Infomax), and data were downsampled to 250 Hz. Trials were segmented into 7-second epochs and manually cleaned based on variance and visual inspection, ensuring no more than 10% trial rejection. Source reconstruction was conducted using individual BEM head models and weighted minimum norm estimation (wMNE), with activity mapped to the MNI template and analysed using the Desikan-Killiany atlas^45^. For a complete description of the experimental design, data acquisition, and preprocessing pipeline, please refer to Corsi et al., 2020^28^.

### 2.2 Preliminary parameters’ selection

The detection of neuronal avalanches involves discretizing the EEG signal, continuous and coarsely sampled, into a series of binary events. This is typically achieved by z-scoring the signal and applying a threshold, with an avalanche defined as a sequence that begins when the signal of at least one brain region exceeds this threshold and ends when all regions’ signals return below the threshold^39^ *(Figure 1b)*. To enable the detection of neuronal avalanches, it is essential to define two key parameters: the z-threshold and the minimum avalanche duration. The z-threshold (θ_av_) determines the signal amplitude above which an event is considered part of a neuronal avalanche. Modifying this value can significantly impact the sensitivity of avalanche detection, thereby influencing the amount and reliability of information extracted from the signal. The second parameter, the minimum avalanche duration (λ_min_av_), sets the lower limit on the temporal extent of an avalanche for it to be considered valid. This criterion ensures that only events with sufficient temporal length are included in the analysis. Careful optimization of these parameters is crucial to ensure that the detected neuronal avalanches reflect meaningful neurophysiological activity rather than artifacts. This process helps minimize the influence of short and transient non-neural artifacts, such as muscle activities or eye-movement-related events, thereby improving the validity of the derived features.

To determine the minimum possible duration of neuronal avalanches, we grounded our selection in established neurophysiological knowledge regarding brain region activation during motor and motor imagery tasks. Specifically, we evaluated three durations of λ_min_av_ (5 ms, 50 ms, and 80 ms) each corresponding to distinct phases of motor-related neural processing. A duration of 5 ms captures the very early phase of sensory processing. This brief interval is associated with fast synaptic activation and marks the onset of sensory or motor preparation^46^. A duration of 50 ms corresponds to the activation of premotor and supplementary motor areas, which are responsible for formulating the motor plan required for movement initiation. This stage reflects the engagement of neural circuits involved in coordinating muscle activity^47^. Finally, a duration of 80 ms reflects the activation of the primary motor cortex (M1), at which point the cortex begins transmitting motor commands to the spinal cord. This marks the transition from planning to motor execution^48^.

For selecting the minimum signal excursion considered relevant the z-threshold θ_av_, we based our parameter choice directly on the signal itself, using the normalized values of the different signals (zero mean and unit variance) as a reference. So, for each subject we evaluate as possible θ_av_ the set of values {μ, μ+ σ, μ+ 2*σ, μ+ 3*σ, μ+ 4*σ, μ+ 5*σ}

In a preliminary analysis, we have evaluated the parameters combinations providing a non-null number of neuronal avalanches and the neuronal physiological validity of their duration (>5ms). Hence, we reduced our possible combinations of θ_av_ and λ_min_av_ to ten possibilities indicated in *Table 1*

**Table 1;.**
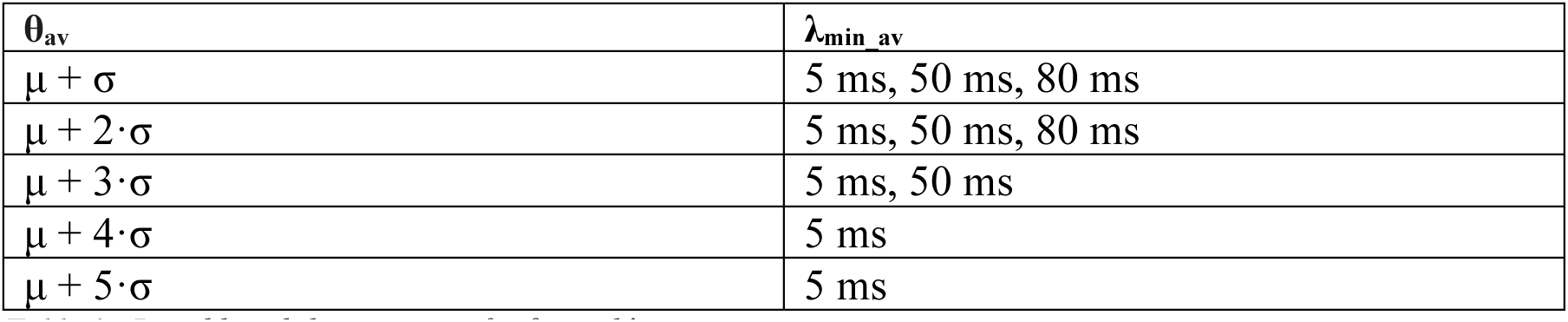
Possible valid parameters for θ_av_ and λ_min_av_.

| $\theta_{av}$ | $\lambda_{min\_av}$ |
| --- | --- |
| $\mu + \sigma$ | 5 ms, 50 ms, 80 ms |
| $\mu + 2\sigma$ | 5 ms, 50 ms, 80 ms |
| $\mu + 3\sigma$ | 5 ms, 50 ms |
| $\mu + 4\sigma$ | 5 ms |
| $\mu + 5\sigma$ | 5 ms |

### 2.3 Features extraction

We analysed binarized avalanche data derived from source-reconstructed EEG signals, mapped onto 68 cortical regions of interest (ROIs) as defined by the Desikan-Killiany atlas^45^. Each neuronal avalanche produced a spatiotemporal activity pattern represented as a binary matrix, where time points exceeding a predefined z-threshold were marked as 1—these instances were considered “activations” *(Figure 1b)*. Neuronal avalanches consist of discrete activations, and the distribution of their durations serves as a key indicator of underlying system dynamics. Guided by this insight, we extracted two primary features to characterize brain dynamics: avalanche length (λ_av_) and activation count (α_av_). Avalanche length (λ_av_) captures the temporal extent of neural propagation. To investigate how this parameter evolves over the course of training, we computed the average duration of avalanches within each trial, followed by averaging across trials for each subject. The second feature, activations count (α_av_), the sum of time samples over threshold θ_av_ across all the ROIs, was used to assess cortical engagement during both motor imagery and rest conditions. For each trial, k, we quantified the total number of activations across all avalanches. To account for variability in avalanche duration, the activation count for each avalanche was weighted by its temporal length. The overall weighted activation score for a given trial was calculated by dividing the total number of weighted activations (across all time samples and ROIs) by the cumulative duration (in time samples) of all avalanches within the k^th^ trial,

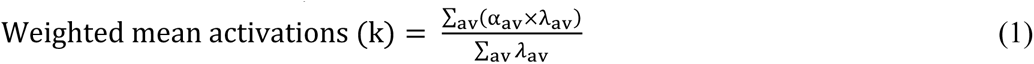

Once computed, these trial-level activation scores were averaged across all trials for each subject and cardinal position, resulting in a single mean activation score per subject.

We extracted these features for each subject, each condition (Rest and MI) and each of the four sessions *(Figure 1c)*.

### 2.4 Statistical analysis

To assess the effects of state (motor imagery vs. rest) and training session (sessions s1 to s4) on the two avalanches’ features extracted across the different possible pairs of parameters θ_av_ and λ_min_av_, we employed a combination of non-parametric statistical approaches, each tailored to different inferential levels: global versus local effect evaluation.

To evaluate the global effects of state, session training effect, and their interaction without relying on normality assumptions, we used a permutation-based approach of ANOVA test^49^ with 10,000 iterations. In each iteration, the dependent variable was randomly shuffled across subjects and conditions, and F-values were recalculated. Empirical p-values were derived by comparing the observed F-statistics to the distribution of permuted values. This approach allows a global assessment of whether there are systematic effects across all sessions and conditions, rather than focusing on individual pairwise comparisons.

To complement the global analysis, we applied non-parametric tests tailored for more localized inference. To assess the learning effect, we conducted the Friedman test^50^ separately for the MI and resting conditions. This allowed us to examine whether significant changes occurred across the four sessions within each condition, capturing temporal effects. To evaluate the task condition effect, we performed the Wilcoxon Signed-Rank Test^51^ independently for each session. This enabled a direct comparison between the MI and resting conditions at each time point, offering a more granular view of condition-specific differences.

### 2.5 Repeated correlation

We implemented a repeated measure correlation analysis to identify which parameters get a significant correlation with BCI-score and use them as possible candidate for our predict model.

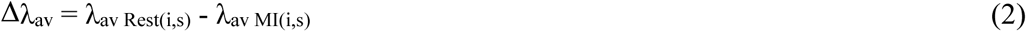

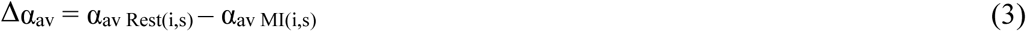

where *i* and *s* index subjects and sessions, respectively. This was repeated for each pair of parameters (θ_av_ and λ_min_av_).

To estimate the correlations between BCI scores and, respectively, the features Δλ_av_ and Δα_av_, we performed repeated-measures correlations^52^ which control for non-independence of observations obtained within each subject without averaging or aggregating data *(Figure 1d)*. It allows us to get a more accurate estimate of the strength of correlation in longitudinal data than standard Pearson correlation.

### 2.6 ROIs selection

To evaluate the effect of number of ROIs, we performed ROI selection to determine whether this approach reduces noise in the dataset, thereby leading to higher correlation and improved prediction accuracy.

Following the computation of activation values across trials, we derived ROI-level features to analyse spatial patterns of cortical engagement during the BCI training. Each trial was represented as a matrix of binarized reconstructed data. For each trial, the activity within each ROI was aggregated and weighted by segment duration, resulting in a length-weighted mean activation per ROI. To reduce inter-subject variability and enable comparisons across sessions and conditions, ROI activations were normalized per subject. Specifically, for each subject and ROI, activations were scaled as a percentage of the maximum activation observed during the first Rest session:

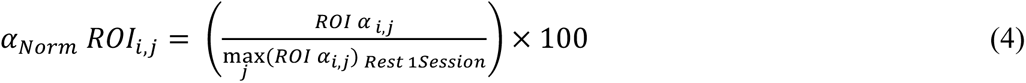

where *i* and *j* denote the subject and ROI index, respectively.

Importantly, while we used the maximum as the reference in this study, the normalization could equivalently be based on the mean, median, or minimum ROI activation without substantially altering the outcome. Since all measures were applied within-subject and per ROI, the resulting normalized values preserved relative spatial patterns and trends across conditions, regardless of the specific scaling baseline. After computing activation values at the single-ROI level for each subject, session, and condition (MI and resting), we performed within-subject paired t-tests to compare the two states independently for each session. This step allowed us to identify ROIs showing the greatest differences in activation between the two conditions. The resulting t-value maps—one per subject and session—were then used as input for a two-way ANOVA, to jointly consider task condition effect and learning effect across subjects. This process ultimately yielded a set of significant ROIs that were robust across both individuals and training sessions.

### 2.7 Predictive models

To investigate to which extent the extracted features could be relevant to predict the BCI score in the subsequent session we implemented two models: a regression model to estimate the exact BCI score and a classification model to determine whether the score indicates effective BCI control *(Figure 1d)*. Both models take the same inputs — Δλ_av_ and Δα_av_ of the first three sessions — and on both a grid search for hyperparameters is done.

To predict the BCI-score at fourth session we implemented a Longitudinal Support Vector Regression model, inspired by the framework of Du et al., 2015^53^. Within this approach, we assumed repeated measurements for each subject, i, over multiple time points, represented as a list of subject-specific matrices 𝑋_*i*_∈ 𝑅^*s*×*f*^ and outcome vectors 𝑦_*s*_∈ 𝑅^*s*^, where *s* is the number of sessions and *f* denotes the number of features. To model temporal trends, we introduced a temporal weight vector 𝛽 ∈ 𝑅^*s*^ that projects both predictors and outcomes into a shared subspace representing longitudinal progression. The regression problem is formulated in the dual space using a quadratic programming (QP) framework, where the objective includes an ε-insensitive loss and dual variables γ∗,γ associated with each subject. The Gram matrix G encodes temporal covariance using:

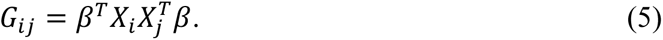

After solving the QP to obtain the dual coefficients α=[γ∗, γ], we estimated the optimal temporal trend vector β by solving a linear system derived from the dual variables and the outcome projections. This iterative estimation allows joint modelling of within-subject dependencies and between-subject variation. Prediction for new subjects is performed by computing a weighted combination of inner products between the new subject’s data and those of the support vectors, modulated by the learned β and dual coefficients.

To predict the capability to control a BCI device at fourth session we implemented a Longitudinal Support Vector Classifier (LSVC), extending the standard SVC to account for repeated measurements across sessions, as proposed by Chen and Bowman, 2011^54^. The aim is to jointly estimates the separating hyperplane parameters and the temporal trend parameters using quadratic programming, so a key challenge that we address is how to jointly estimate the parameter vectors β and α. In this framework, longitudinal data are collected from N subjects over S sessions, with features f measured at each session point. For each subject, the data form a matrix 𝑋_s_∈ 𝑅^S×f^, and the classification label is y_s_∈{0,1}. To incorporate the temporal structure, we model each subject’s data as a weighted combination of measurements across time:

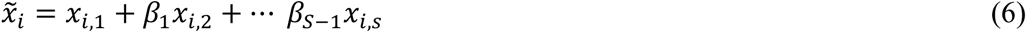

where 𝛽 = (1, 𝛽_1_, 𝛽_2_ … 𝛽_s-1;_) encodes the temporal trend.

The LSVC algorithm operates through an alternating optimization process. It begins by solving the dual SVM problem via quadratic programming, using the current value of β. In this step, the kernel matrix 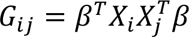 captures temporal dependencies through β. Following this, β is updated by solving a regularized linear system, which is derived from the current dual coefficients and the subject data. This iterative procedure continues until convergence. Here, the threshold to distinguish subjects able/unable to control a BCI-device was set at 57%, which represents the chance level in our dataset^55^.

To evaluate the model performance, we employed leave-one-out cross-validation (LOO), a rigorous validation method particularly well-suited for small datasets. In each iteration, one subject’s data was reserved as the test set, while the data from the remaining subjects were used to train the model. This process was repeated until every subject had served once as the test case, ensuring comprehensive evaluation across the entire dataset.

To assess the effectiveness of our models, we compared their performance against different baselines. The first was a conventional implementation of Support Vector Regression (SVR) and a Support Vector Classifier (SVC), which do not incorporate temporal modelling. The second group of models involved applying our longitudinal model to data in which the session order had been randomly shuffled. This allowed us to isolate and quantify the specific contribution of temporal learning effects, providing a benchmark referred to as the random sessions condition.

## Results

### 3.1 Changes of avalanche lengths and activations across time

At a global level, as assessed using a permutation ANOVA *(Figure 2)*, no significant effects were observed for either condition (Rest vs. Motor Imagery) or training (across sessions) in any combination of parameters (θ_av_ and λ_min_av_). This finding holds for both avalanches’ features: length (λ_av_) and activations count (α_av_). However, a more granular analysis of λ_av_ and α_av_ revealed, for some parameters, a significant difference (*p* < 0.048) between the resting and MI conditions during the fourth session. In addition, a significant learning effect (*p* < 0.034) across sessions within the MI condition were also observed.

**Figure 2:**
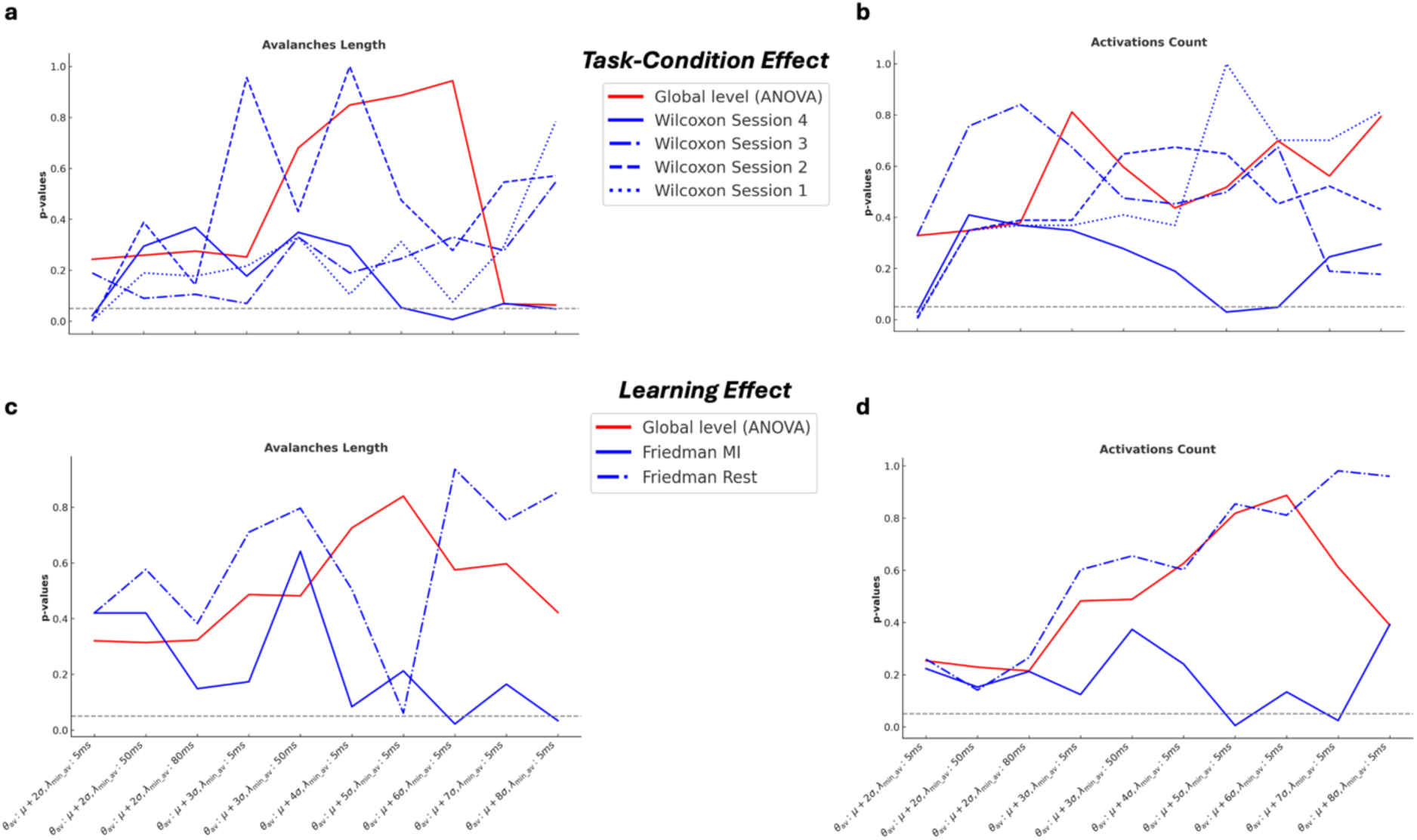
Task and learning effects on different parameters: **a)** Task-condition effect on avalanches length; **b)** task-condition effect on activations count; **c)** Learning effect on avalanches length; **d)** Learning effect on activations count. Red lines correspond to the ANOVA general level while the blues lines correspond to the local effects results from Wilcoxon and Friedman test. Grey dashed line represent the p-values significative threshold at 0.05.

**Figure 2:**
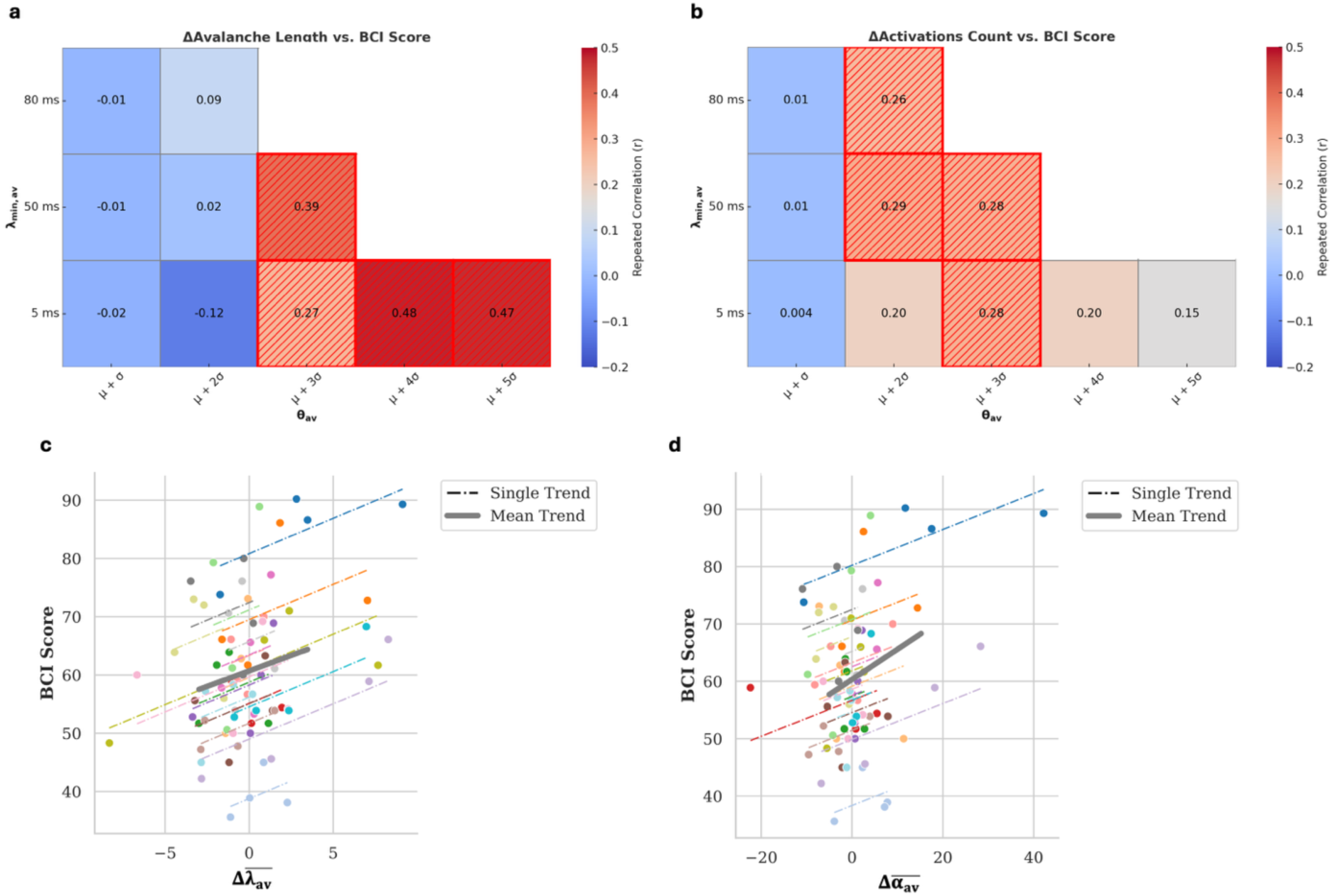
Repeated correlation and trends across different sessions between BCI score and features changes. **a)** mean difference of avalanches’ length (Δλ_av_)and **b)** mean difference activation (Δα_av_) across all tested pairs of parameters (θ_av_, λ_min_av_). Significant correlations (p<0.05) are highlighted using different textures. **c)** repeated correlations trend across different sessions between BCI-score and Δλ_av_ and **d)** between Δα_av_ and BCI-score. Each coloured dashed line corresponds to one subject while the grey bold line identifies the trend across all the subjects. The pairs of parameters (θ_av_, λ_min_av_) used in **c)** and **d)** were those that achieve the best prediction performance: θ_av_: μ+3σ, λ_min_av_ : 50ms

All results reported here were derived using the broad frequency band [8–35 Hz]. We additionally examined other frequency bands, including the alpha band [8–12 Hz], beta band [12–30 Hz], low gamma band [30–50 Hz], and theta band [3–8 Hz] (analyses not shown here). In the alpha and beta bands, the results generally mirrored the main patterns observed in the broad frequency band, though with fewer significant parameter pairs. This suggests that the broader frequency range offers a more holistic and integrative perspective across our analyses. This observation is consistent with previous studies indicating that arrhythmic neural activity can contribute to broadband EEG signals^34^. Consequently, these findings underscore the importance of broadband analyses in our study.

### 3.2 Repeated Correlation

Using the same combinations of parameters, we also observed significant correlations between the changes in avalanche length and activations with the BCI performance scores (*Figure 3a-b*). Specifically, significant positive correlations (*p* < 0.033 for Δλ_av_ and *p* < 0.040 for Δα_av,_ respectively) were found in different conditions *(Supplementary Materials Table 3)*.

**Figure 3:**
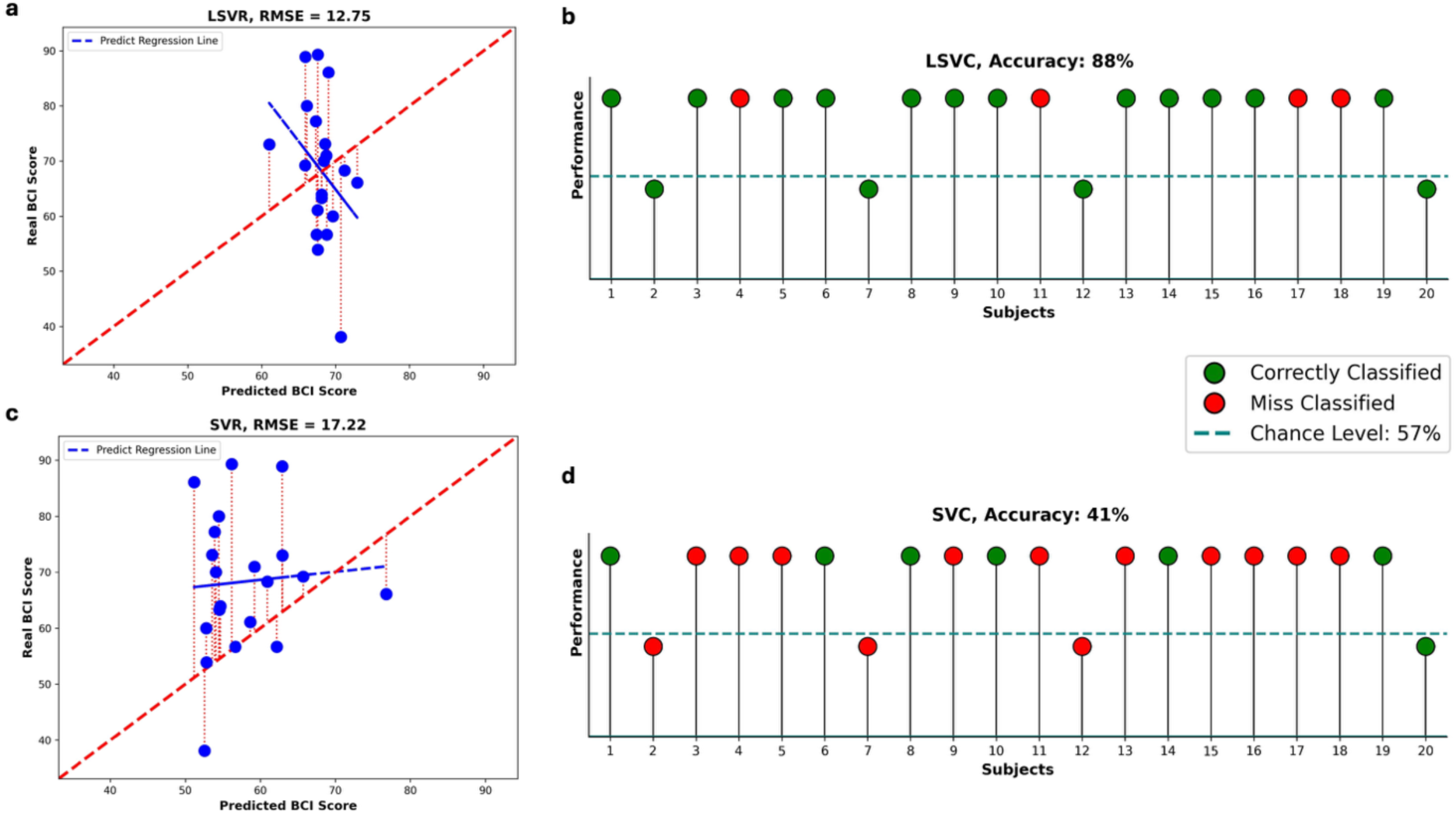
Prediction results. **a)** Predicted BCI-score from a longitudinal regression model, compared to a **c)** standard SVR model. Each point represents one subject. The red dashed lines indicate the prediction error between actual and predicted values. The bold red line shows the optimal prediction trend, while blue lines represent the predicted trends. **b)** Prediction of control results using a longitudinal predictive model, compared to a **d)** standard SVC model. Each ball represents one subject. Subjects above the dashed line are considered able to control the outcome. Green balls indicate correct predictions of control ability, while red balls represent misclassifications.

In all cases, the correlations were positive, indicating that as the difference between resting and MI conditions increased across sessions, the BCI performance also improved. Notably, this progressive difference was associated with a monotonic increase in the strength of the correlation between both Δλ_av_ and Δα_av_ with BCI scores, which was observed in most participants *(one example in Figure 3c and Figure 3d)*.

### 3.3 Prediction results

Among the possible coupled parameters that show a significant correlation between the change of our features and BCI-score, the highest predictive performance is achieved for the parameters θ_av_= μ + 3σ and λ_min_av_ ≈ 50ms. Using the pair of parameters that yielded significant correlations with BCI performance, we implemented two longitudinal predictive models: a regression model (Longitudinal Support Vector Regression, LSVR) and a classification model (Longitudinal Support Vector Classifier, LSVC). These models were designed to predict each subject’s BCI score one session ahead based on data acquired up to the previous session.

#### BCI performance prediction

The LSVR model generates a continuous predicted BCI score, which was evaluated using the Root Mean Square Error (*RMSE*) relative to the actual continuous BCI scores. Overall, the predictions produced by the LSVR model exhibit a lower error compared to those obtained using the standard SVR model (*RMSE_LSVR_* = 12.7481; *RMSE_SVR_*= 17.2186) *(Figure 4 a-c)*. In contrast, the difference in performance between the LSVR model and the session-shuffled variant is less pronounced (*RMSE_LSVR_*= 12.7481; *RMSE_LSVR_* with random session shuffling = 13.3571) *(Supplementary Materials, Figure 1)*.

#### BCI control effectiveness prediction

Our primary objective was to predict whether a subject will be able to control the BCI system in the following session—based on individual learning progress—or whether additional training would be necessary. For this reason, we also implemented the LSVC model to perform a binary classification. We defined successful control using a performance threshold set at the chance level of 57%, as reported by Müller-Putz et al. 2008^55^ Thanks to LSVC, we achieved 88% accuracy in predicted performance. Four subjects on twenty total were misclassified *(Supplementary Materials Figure 1c)*. LSVC model significantly outperformed the standard SVC, achieving 88% compared to 41% accuracy *(Figure 4 b-d)*. Additionally, to evaluate the influence of the learning effect over time, we performed a session-shuffling control analysis. When the temporal order of the three sessions was randomly shuffled, classification accuracy dropped to 47% *(Supplementary Materials Figure 1).* These results suggest that the longitudinal structure of the data contributes meaningfully to classification performance, although the limited number of sessions of our study (three) may constrain the observable effect.

### 3.4 ROIs selection effect

Following the ROI selection procedure outlined in the Materials and Methods section, we identified a consistent set of significant ROIs across all subjects for each parameter combination. This set *(Supplementary Materials Figure 2)* captures both condition-related differences (Rest vs. MI), as indicated by the t-values, and learning effects over time, as revealed by ANOVA across sessions. Different combinations of θ_av_ and λ_min_av_ consistently identified ROIs predominantly located in areas associated with higher-order cognitive functions, including decision-making, attention regulation, and motor planning. These include the right inferior parietal lobule, right isthmus cingulate, and left precuneus, as well as occipital regions in the visual cortex directly involved in visual feedback stimulation.

It is worth noting that the parameter combination using a θ_av_ = *μ + σ* did not yield any significant ROIs. Therefore, in this section, we focus exclusively on parameter combinations with larger z-thresholds. For each selected set of significant ROIs, we extracted the same two features λ_av_ and α_av_. Using only these selected ROIs, we were able to detect both local and global statistical effects *(Supplementary Materials Table 4, Table 5)*.

Locally, we observed significant differences between Rest and MI conditions (*Figures 5a-b*) during the fourth session, for both features: λ_av_ and α_av_ (Wilcoxon test, *p* < 0.036, *p* < 0.030, respectively). Additionally (*Figures 5c-d*), a significant learning effect was present within the MI condition across sessions (Friedman test, *p* < 0.021and *p* < 0.042 for λ_av_ and α_av,_ respectively). Globally, using a 10.000-permutations ANOVA, we also observed significant task-condition (*p* < 0.036 and *p* = 0.045 for λ_av_ and α_av,_ respectively) and learning effects (*p* < 0.024) for λ_av_ *(Supplementary Materials Table 4, Table 5)*.

**Figure 5:**
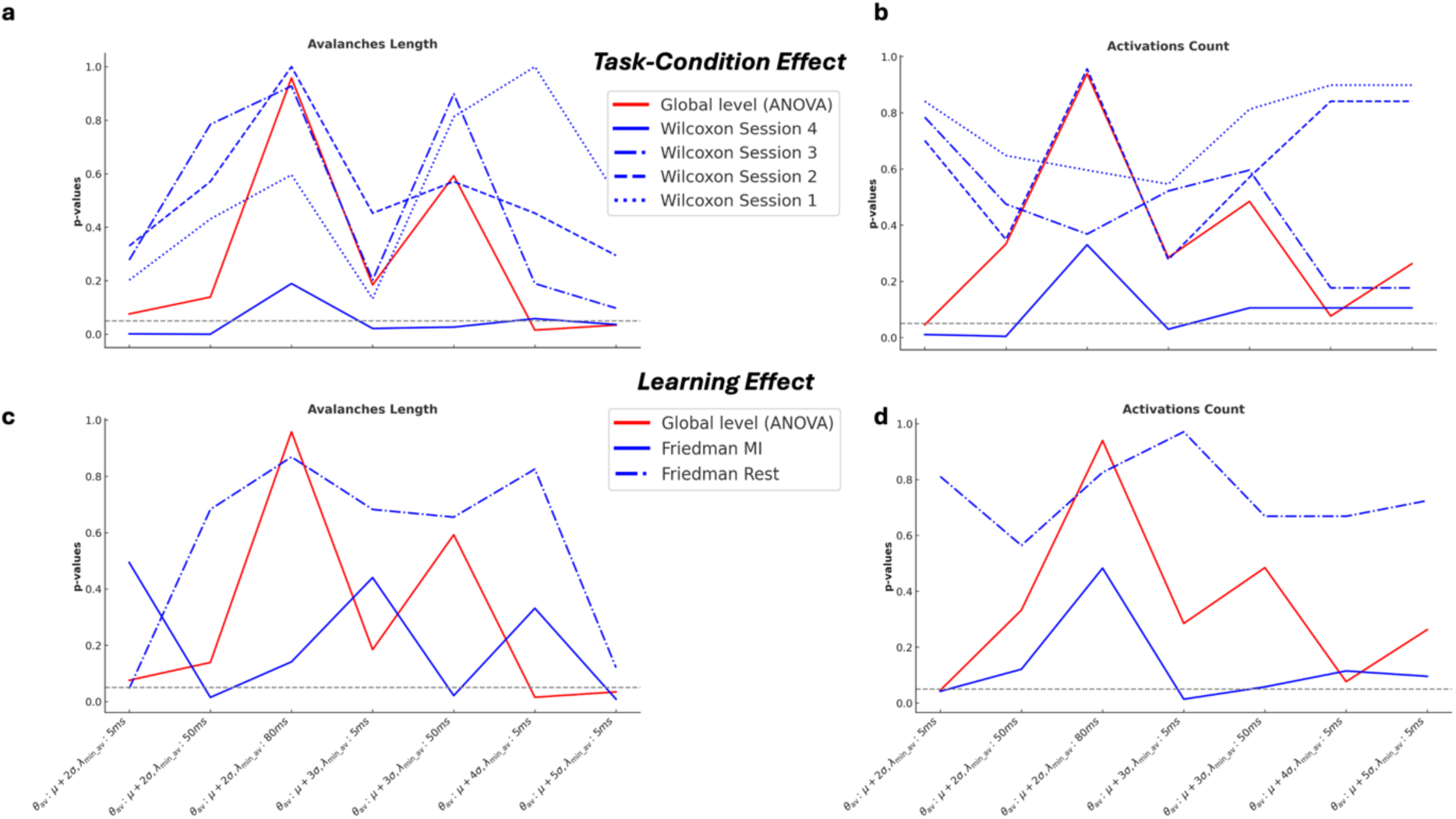
Task and learning effects on different parameters. **a)** Task-condition effect on avalanches length; **b)** task-condition effect on activations count; **c)** Learning effect on avalanches length; **d)** Learning effect on activations count. Same stipulations as in the caption of Figure 2.

Focusing on avalanches’ length distributions, we observed a progressive, condition-specific modulation of neural dynamics over the course of the BCI training sessions *(Figure 6)*. Specifically, during MI condition, there was a consistent increase in average avalanche length across sessions, evidenced by a rightward shift and sharpening of the corresponding probability density distributions (PDFs). In contrast, the Rest condition exhibited no significant change, with its PDFs remaining largely stable across sessions. At the first session, MI and Rest distributions overlapped, but diverged progressively with training, this separation of the distributions reflects a dynamic learning process. By session four, the resting distribution had become broader and flatter, suggesting increased inter-subject variability, whereas the MI distribution becomes narrower and more peaked, reflecting a more consistent and uniform neural response among participants. Interestingly, the MI distribution also developed a bimodal shape, revealing two distinct participant subgroups: one displaying significantly longer avalanche lengths indicative of MI proficiency, and another whose dynamics remained comparable to Rest. This divergence underscores individual differences in the ability to develop effective motor imagery strategies. Notably, this pattern was consistent both with and without applying spatial ROI selection.

**Figure 6:**
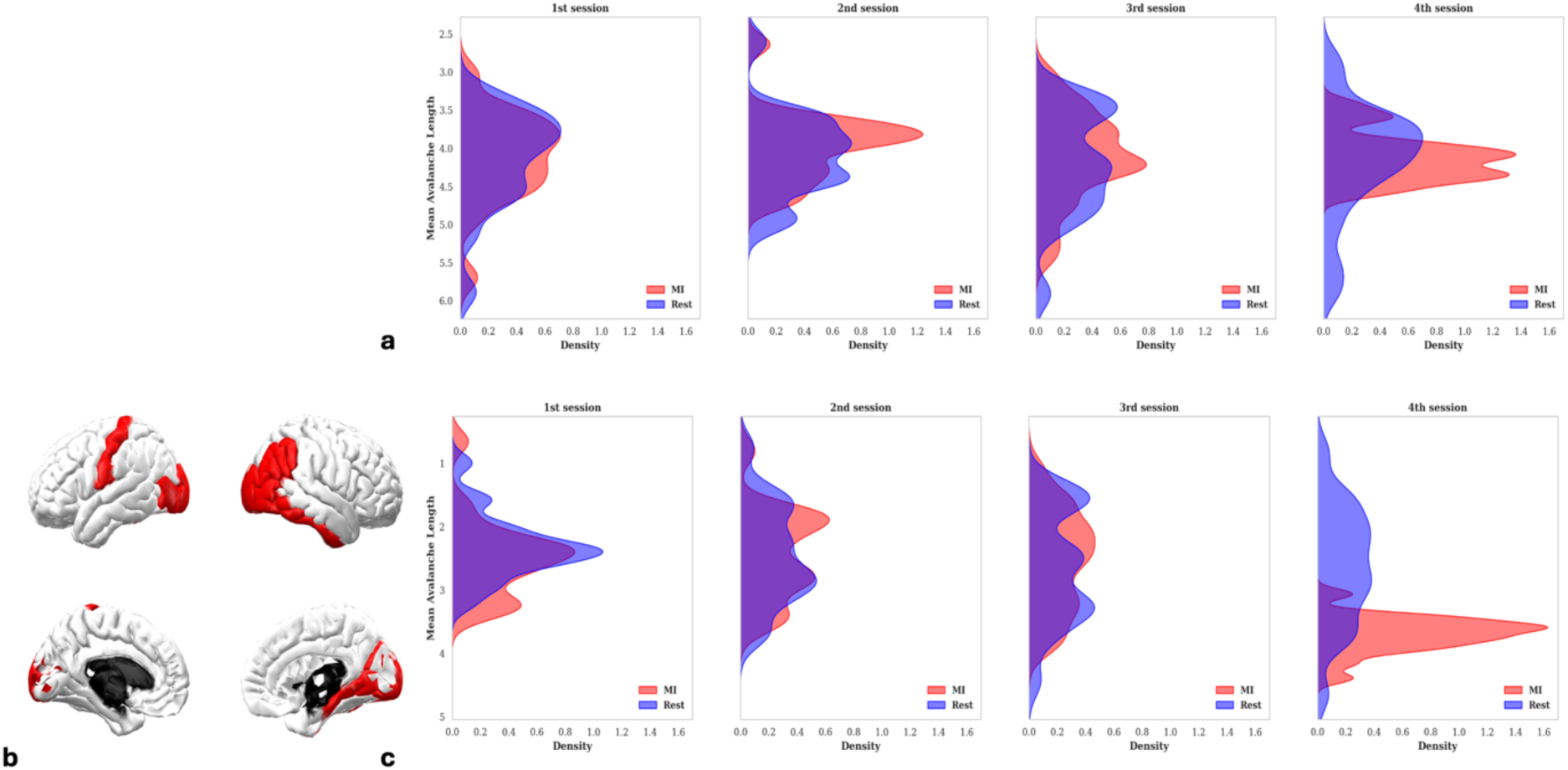
Evolution of mean avalanche length. Probability distribution analysis across all the subjects repeated for all the training sessions (from the first session on the left to the fourth session on the right) to represent the different distributions in MI task condition (red) and in resting state (blue). **a)** over the entire dataset and **c)** over a selected set of ROIs. The selected ROIs are shown in **b.**

Repeated correlation analyses between BCI performance and Δλ_av_, Δα_av_ confirmed our earlier results, revealing significant associations between both features and BCI scores (*p* < 0.047 & *p* < 0.046, for Δλ_av_ and Δα_av_, respectively).

As in the analysis with all ROIs, we trained two longitudinal predictive models—LSVR (regression) and LSVC (classification)—using selected ROIs and the parameter combinations that showed significant repeated correlations. Best results are obtained using the parameter setting of θ_av_ *= μ + 2σ* and *λ_min_av_ = 50ms* for both Δλ_av_ and Δα_av_ (*Supplementary Materials Figure 3c-3d)*. The LSVR model yielded a slightly lower RMSE (10.3931) compared to the model trained on the full dataset without ROI selection (RMSE = 12.7481). Furthermore, the regression trend produced by the LSVR model more closely aligned with the observed data following ROI selection *(Figure 7a)*. To assess the benefit of incorporating longitudinal information, we compared the LSVR model with the standard SVR model that does not account for session order. LSVR achieved a significantly lower *RMSE* (10.3931 vs. 17.0468) *(Figure 7a-7c)*, highlighting the added value of modelling session progression. Additionally, the sessions random shuffle leads to a slight increase in the *RMSE* value (10.3931 vs. 13.3219) *(Supplementary Materials Figure 4)*, the results still suggest a meaningful contribution of longitudinal structure.

**Figure 7:**
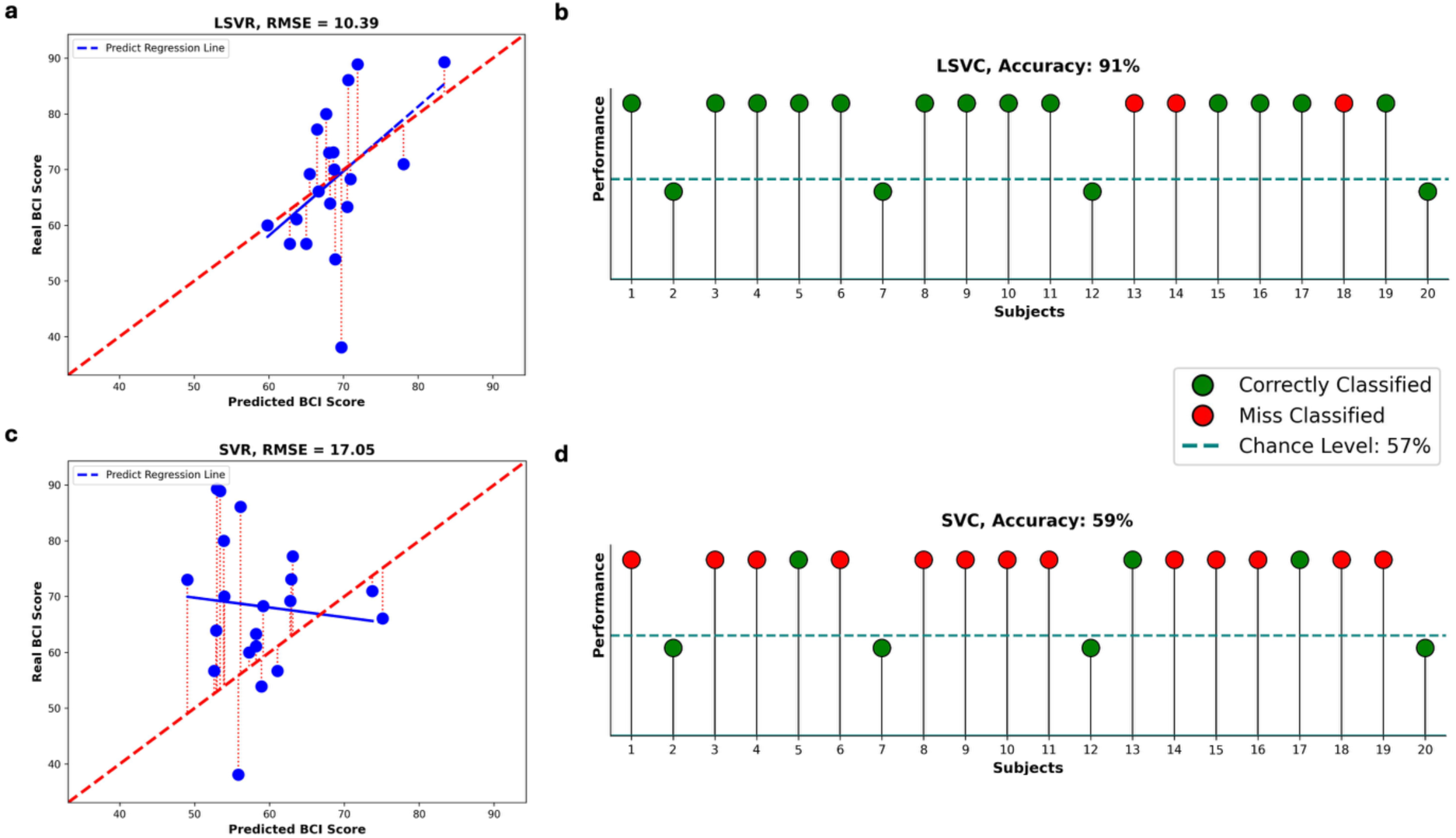
Prediction results over a selected set of ROIs. **a)** Predicted BCI-score from a longitudinal regression model, compared to a **c)** standard SVR model. Each point represents one subject. The red dashed lines indicate the prediction error between actual and predicted values. The bold red line shows the optimal prediction trend, while blue lines represent the predicted trends. **b)** Prediction of control results using a longitudinal predictive model, compared to a **d)** standard SVC model. Each ball represents one subject. Same stipulations as in the caption of Figure 4.

The LSVC model achieved an overall classification accuracy of 91%, with only three false negatives out of 20 subjects *(Supplementary Materials Figure 4c)*, demonstrating its effectiveness in distinguishing successful from unsuccessful BCI learners. Compared to the standard SVC model, which does not account for session order, the LSVC model demonstrated substantially superior performance, achieving an accuracy of 91% versus 59% *(Figure 7b-7d)*. To further examine the role of temporal structure, we performed a random shuffle of session order. This manipulation resulted in a decrease in classification accuracy to 84% *(Supplementary Materials, Figure 4)*, indicating that the inclusion of longitudinal information contributes meaningfully to model performance. Nevertheless, given the limited number of sessions (n = 3), the full magnitude of this effect may be underestimated.

## Discussion

This study aimed to investigate the neural mechanisms underlying motor imagery-based brain– computer interface training, with a particular focus on identifying condition-induced neural modulations and predictors of individual performance. BCI and neurofeedback outcomes are shaped by a complex interplay of both stable and dynamic factors, including psychological traits, cognitive abilities, and neurophysiological patterns. Collectively, previous findings highlighted that successful BCI learning depends not only on localized brain activations but also on a broader reorganization and downregulation of cognitively demanding network interactions. Building on this knowledge, the present study introduces a functional connectivity approach designed to also capture the spatiotemporal behaviour of aperiodic neural signals with the final goal to design personalized training programs.

### 4.1 Task Condition Effect and Learning Effect

Specifically, we characterized cortical dynamics by two avalanches’ features (the length and activation count) and found that the effect of task condition (Resting state vs. Motor Imagery) became most pronounced during the last and fourth session. Additionally, a more robust learning effect emerged across sessions within the MI condition, as compared to rest. Taken together, these results point to an underlying learning process in which the brain progressively adapts to enhance attention toward the imagined right-hand grasping task. This adaptation amplifies the separation between resting and motor imagery states, reflecting functional plasticity developed through MI-BCI training. Additional support for this interpretation comes from the probability distributions of avalanche lengths *(Figure 6)*, which further illustrate how neural dynamics evolve with practice. Overall, these observations are consistent with established learning effects in MI-based BCI training, where neural activity becomes increasingly organized and distinct from rest as participants gain familiarity with the task^56,57^.

Our results suggest that applying a ROI selection is beneficial, as it does not compromise the statistical significance of our findings but, rather, enhances our ability to detect significant differences between MI and resting conditions across all sessions, as well as robust learning effects in both brain states. One should keep in mind that the ROI selection method we put forward generates a distinct set of ROIs for each parameter combination and occasionally includes regions outside the primary motor areas targeted by the BCI. However, it is important to note that these non-motor areas, including associative and visual processing regions, are likely to be involved in the training process, contributing to motor imagery through integrative and supportive cognitive mechanisms. This interpretation is further supported by the experimental design, in which subjects receive three seconds of visual feedback during each trial, likely engaging visual and associative cortices alongside motor areas^28^. These results highlight a key insight: not all ROIs contribute meaningful information. By excluding non-informative ROIs, we enhanced the overall signal-to-noise ratio, enabling us to uncover consistent, global differences between resting and MI states across sessions, as well as clear learning-related effects. These observations are in line with the findings of Mencel et al., 2022^58^, who found that electrode location influenced both the magnitude and timing of the ERP responses, and highlighting the spatial specificity of training-induced cortical changes, supporting the hypothesis of distributed neuroplastic adaptations.

To validate the effectiveness of the proposed features (λ_av_ and α_av_) we further analysed data from the final session by categorizing each trial for each subject as either a successful (Hit) or an unsuccessful BCI control (Miss). For several parameter combinations, both features reliably distinguished between MI Hit and MI Miss trials, Rest Hit and Rest Miss trials, as well as MI Hit versus Rest Hit trials, with statistical significance (always *p* < 0.05, using specific paired parameters *p* < 0.005). Moreover, for most parameter settings that showed strong and consistent correlations with BCI performance (as discussed in the previous section), we observed significant interaction effects (permutation ANOVA, *p* < 0.05). In the case of avalanche length, there was also a significant main effect of task condition (permutation ANOVA, *p* < 0.05). This capacity for consistent reorganization between successful and unsuccessful trials is in alignment with previous results by Corsi et al., 2024^42^. We hypothesized that the spatiotemporal organization of avalanches could, in the future, serve as a novel source of features to support and extend our current findings—offering new, yet unexplored insights into brain activity in BCI applications. These findings strongly support the conclusion that both neuronal avalanche length and activation are directly associated with distinct brain states— Rest and Motor Imagery—and are sensitive to the subject’s ability to successfully perform the assigned BCI task. In other words, these features not only capture brain state differences but also reflect task performance at the trial level *(Supplementary Materials Figure 5)*. An additional and promising insight emerged from comparing ROI occurrence across all trials versus only successful trials. This analysis, which underlies our ROI selection process, revealed a marked increase in t-values activations (Rest vs MI) within motor-related ROIs using only successful trials. This supports the assumption that increased occurrence in motor areas is linked to successful motor imagery performance, thereby validating the rationale behind our ROI selection strategy *(see Supplementary Materials Figure 6)*.

### 4.2 BCI Predictors

Building on this foundation, we investigated the relationship between specific neural features and BCI performance across sessions. Using repeated-measures correlation, we found that changes in avalanche length and activation—specifically the difference between motor imagery (MI) and resting-state conditions (Δ = MI – Rest)—were significantly correlated with individual BCI scores. While feature values during the Rest condition remained stable over time, those in the MI condition were progressively modified. This led to a shift in Δ values from negative (Rest > MI) to positive (MI > Rest), a change that was positively associated with improved BCI control. These findings were further supported by PDF analyses (Figure 6), which showed that higher MI feature values consistently corresponded to better BCI scores, both with and without ROI selection. In the study of Trocellier et al., 2024^59^, authors assessed different neurophysiological predictors on the same dataset, but their analysis was limited to single session runs. In contrast, our results are based on data spanning four sessions, underscoring the robustness of our findings over time.

### 4.3 Generalized Parameters Optimization

Although differences and specific analytical strategies led to some variability in the optimal parameter combinations, our analyses identified a subset of parameter settings that consistently enabled robust detection of neuronal avalanches across subjects, features, and sessions. Among the various z-threshold and minimum avalanche duration pairings tested, we found that several combinations effectively yielded avalanche features (i.e., avalanche length and activations count) that were not only sensitive to task-related brain dynamics, but also significantly correlated with and predictive of BCI performance. We observed converging evidence supporting the use of z-thresholds set at the mean plus 2 to 3 standard deviations and minimum avalanche durations of 5 or 50 ms. These settings strike a balance between physiological interpretability and predictive utility, yielding avalanche dynamics that reflect behaviourally meaningful brain state changes over time. While we acknowledge the nuanced distinctions between feature-specific optima, we propose this parameter range as a general and reliable guideline for applying neuronal avalanche analysis within the BCI context.

### 4.4 BCI and Effective Control Prediction

Despite advances in this area, many existing BCI predictive models rely on single-session data, which fails to capture the evolving dynamics of learning. Some approaches have included stimulation-based predictors, such as transcranial magnetic stimulation^60^ and median nerve stimulation^61^, but these are often limited to short-term or one-off assessments. A notable exception is the work by Ma et al., 2022^62^, who used a multi-day EEG dataset to predict MI-BCI performance using both MI and resting-state data. However, their models were trained exclusively on first-session data, limiting their ability to account for longitudinal learning effects. The objectives of our study align with those of Kübler et al., 2004^22^, who sought to predict the number of training sessions required for individuals—both healthy participants and those with amyotrophic lateral sclerosis (ALS)—to gain effective control over a communication-based BCI system. We notice, however, that their analysis was limited to correlations between discrete sessions and did not account for the dynamic progression of learning across successive sessions.

In contrast, our study demonstrates that robust and reliable prediction of BCI performance is achievable using data from as few as three sessions. By incorporating both resting-state and task-related neural activities, and applying repeated-measures correlation, we can predict fourth-session performance with high accuracy. This approach provides a more dynamic and longitudinal understanding of user learning, representing a significant step forward in adaptive BCI modelling. Given the presence of significant repeated correlations using the same parameter combinations that also yielded robust task condition and learning effects, we took an additional step beyond correlation analysis: we employed our features to predict BCI scores for each individual subject. From a clinical perspective, the ability to forecast BCI performance one session in advance would be highly valuable. This predictive capability is possible because our features encode both brain state discrimination (Rest vs. MI) and learning-related changes over time. To simultaneously account for these two aspects, we used longitudinal models— specifically, Longitudinal Support Vector Regression (LSVR) and Longitudinal Support Vector Classification (LSVC). These models significantly outperformed their non-longitudinal counterparts (SVR and SVC), demonstrating the advantage of incorporating temporal structure into the analysis. Furthermore, results from session-shuffling experiments confirmed the importance of session order, reinforcing the relevance of modelling longitudinal progression. The consistency of the results before and after ROI selection supports our earlier hypothesis: it is feasible to reduce the dataset dimensionality via targeted ROI selection without compromising model performance.

### 4.5 Limitations and Future Directions

To the best of our knowledge, this study is the first to apply longitudinal models in the context of BCI training with the goal of predicting individual BCI performance one session in advance. However, a major limitation lies in the relatively small number of training sessions, modelled as a linear learning trajectory across sessions, and may not accurately capture the complex and variable nature of learning processes in real-world settings. Factors such as physiological fluctuations and environmental influences—both of which can affect daily performance—are therefore not explicitly accounted for in the current model.

Nonetheless, our findings provide a promising framework that could support clinicians in forecasting individual progress and designing personalized training protocols. For some users, few sessions may suffice to gain BCI control, while others may require extended training. Tailoring the number and structure of training sessions to each user could help reduce frustration and improve the likelihood of successful BCI adoption. In future work, we aim to validate our approach using datasets that include a larger number of training sessions. This would allow us not only to improve the prediction of BCI performance one or more sessions ahead, but also to estimate the total number of sessions each subject might need to achieve effective control. The primary objective of our study is to establish a stable, group-level set of regions of interest (ROIs) that remains consistent across subjects while preserving the fundamental criterion of capturing both the distinction between Rest and motor imagery (MI) brain states and their temporal evolution across training sessions. The development of a unified and interpretable ROI framework not only enhances the scientific interpretability of our results following dataset reduction through ROI selection but also reinforces their potential clinical applicability.

## Conclusion

Our findings demonstrate that the spatiotemporal characterization of the spreading of neuronal avalanches is highly effective in distinguishing between resting-state and motor imagery brain activities. Moreover, they can reliably indicate whether a subject successfully performs the assigned task. These results highlight the potential of neuronal avalanches as powerful biomarkers for tracking and supporting the BCI training process. In this study, we introduce two novel features—avalanche length and activation—that offer a richer and more comprehensive representation of brain dynamics compared to traditional BCI features. While each feature demonstrates slightly different sensitivities to specific parameter combinations, avalanche length consistently outperformed in detecting task-related effects, capturing learning dynamics, correlating with BCI performance, and achieving higher prediction accuracy. Finally, this work proposes a promising strategy to address the issue of BCI inefficiency by supporting the development of personalized training programs. Tailoring the training process to individual learning profiles may help clinicians enhance user engagement, reduce frustration, and ultimately increase the number of successful BCI users.

## Ethics approval and consent to participate

All procedures performed in studies involving human participants were in accordance with the ethical standards of the institutional and/or national research committee and with the 1964 Helsinki Declaration and its later amendments or comparable ethical standards. Written consents were obtained from all participants. The study was approved by the ethical committee CPP-IDF-VI of Paris.

## Data and Code Availability

The study dataset has been fully collected and curated. A dedicated data-descriptor paper is currently in preparation; upon its acceptance, the de-identified dataset and full documentation will be deposited in an open repository with a citable DOI and made publicly available. Until release, controlled access may be granted on reasonable request to the corresponding author.

Code used to import data, analyse data, and generate manuscript figures are available on GitHub: https://github.com/CamiMannino/Neuronal-avalanches-as-a-predictive-biomarker-for-guiding-tailored-BCI-training-programs

## Authors contribution

Conceptualization: C.M., P.S. M.C., and M.C.C. Methodology: C.M., P.S. M.C., and M.C.C. Investigation: C.M. Supervision: M.C.C. and M.C. Data collection and curation: M.C.C. Data processing: C.M. Writing—original draft: C.M. and M.C.C. Writing—review and editing: C.M., P.S. M.C., and M.C.C.

## Supplementary Materials

**Supplementary Materials, Table 1.**
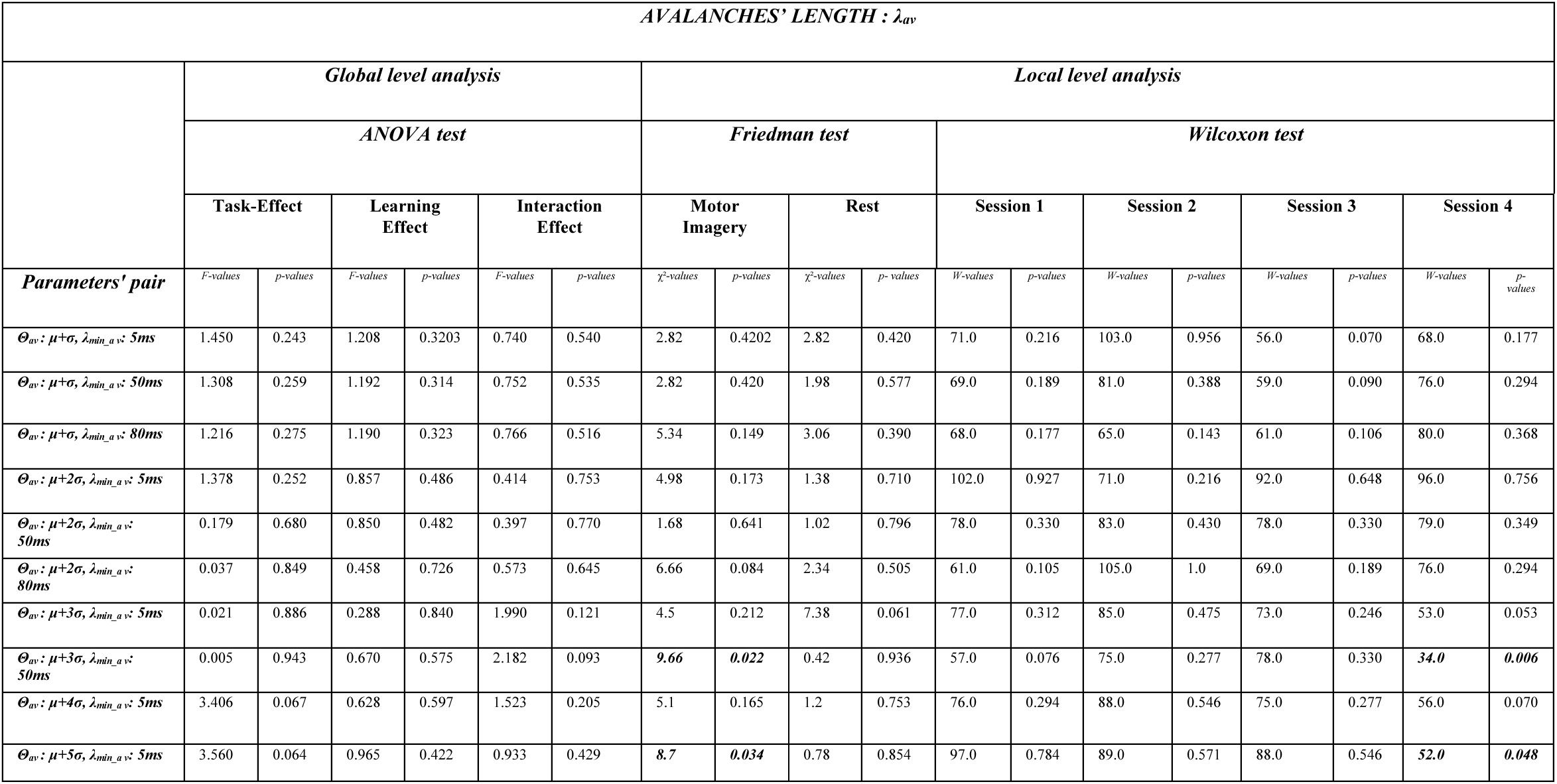
Statistical results of all analysis for Avalanches’ length λ_av_ over all possible parameters’ combinations. In bold the significant values (p < 0.05)

**Supplementary Materials, Table 2.**
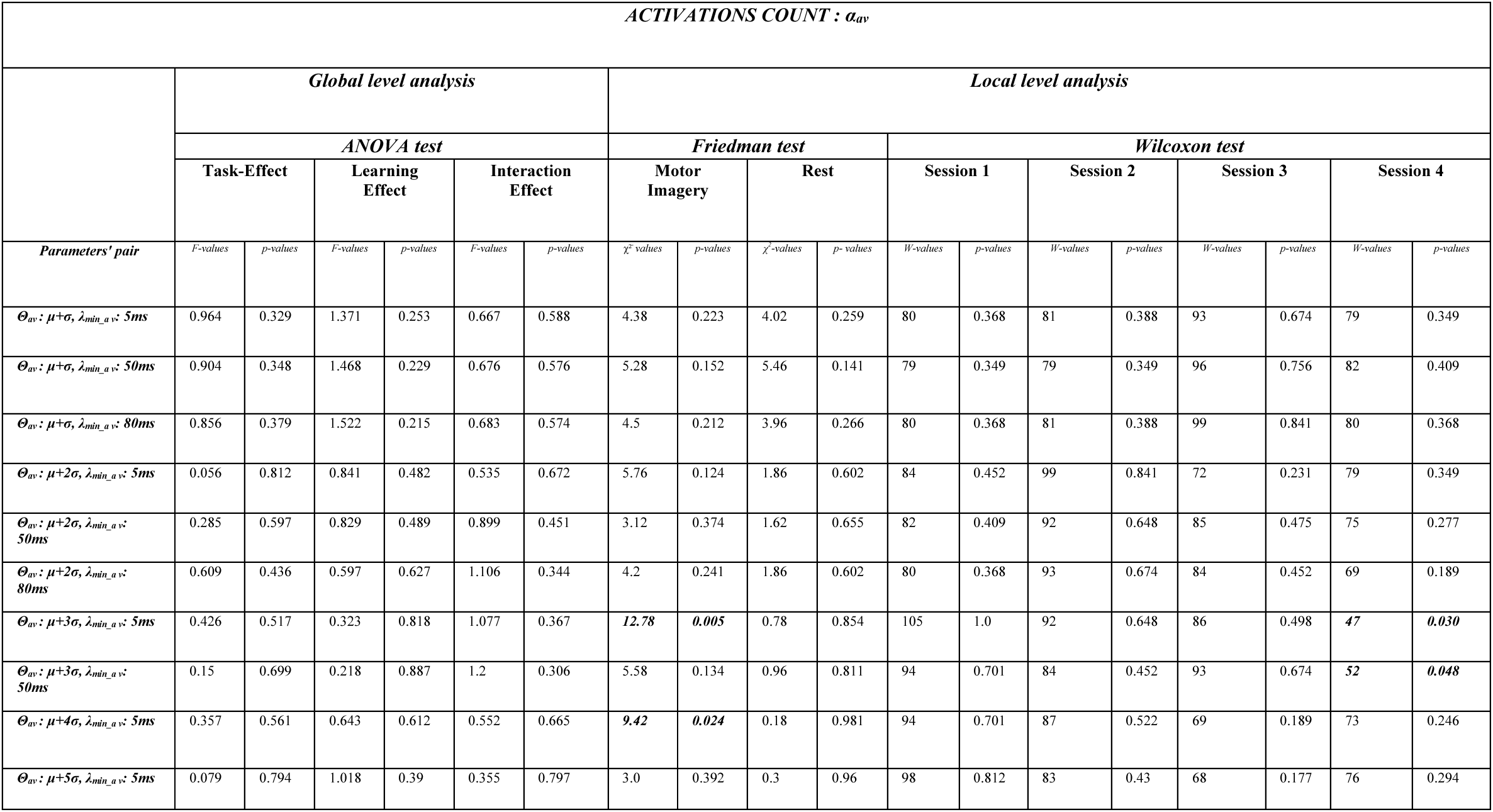
Statistical results of all analysis for Activations count α_av_ over all possible parameters’ combinations. In bold the significant values (p < 0.05)

**Supplementary Materials, Table 3.**
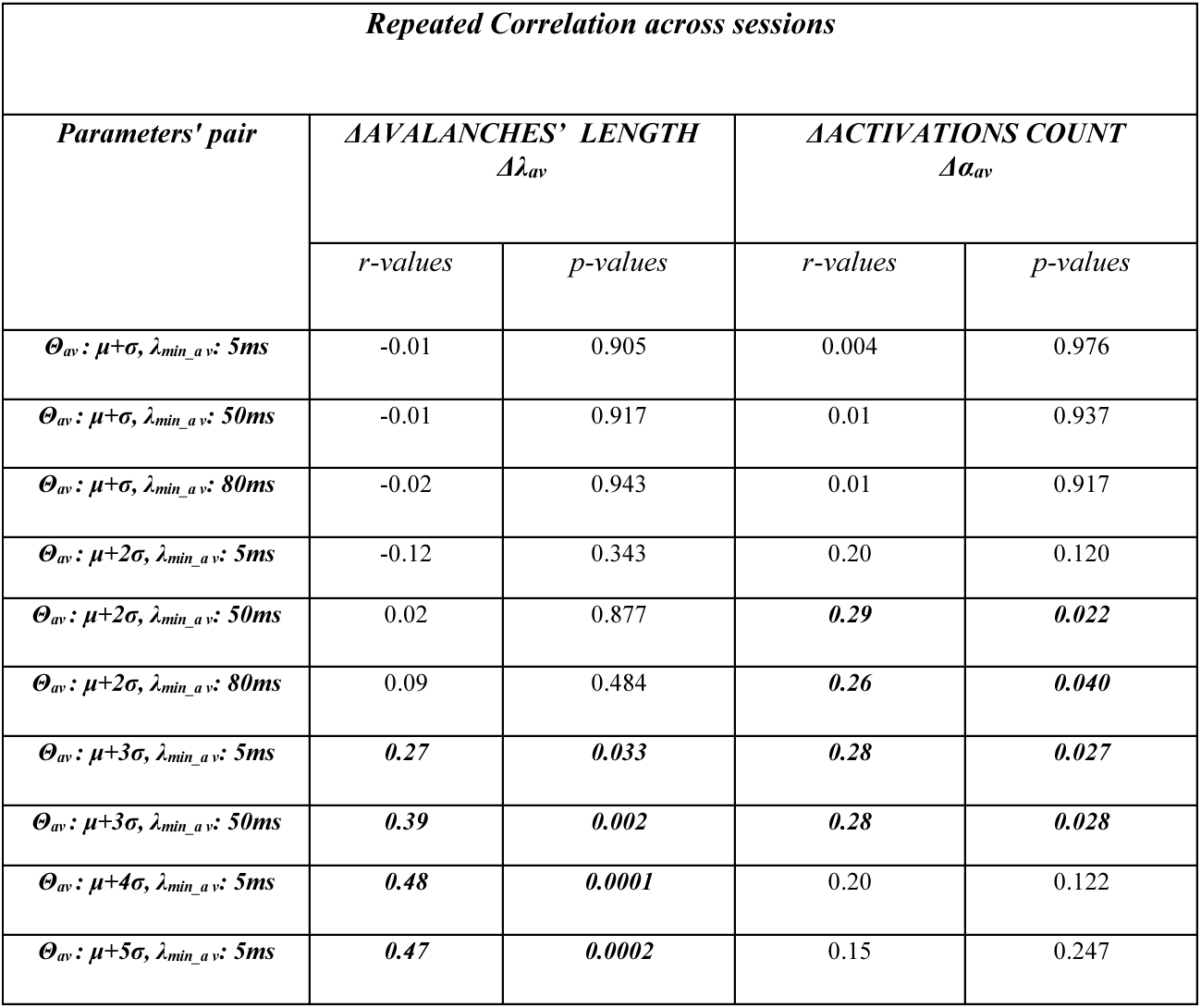
Repeated Correlation results across sessions between both features (Δλ_av_, Δα_av_) and BCI-scores over all possible parameters’ combinations. In bold significant correlations (p < 0.05)

**Supplementary Materials, Figure 1.**
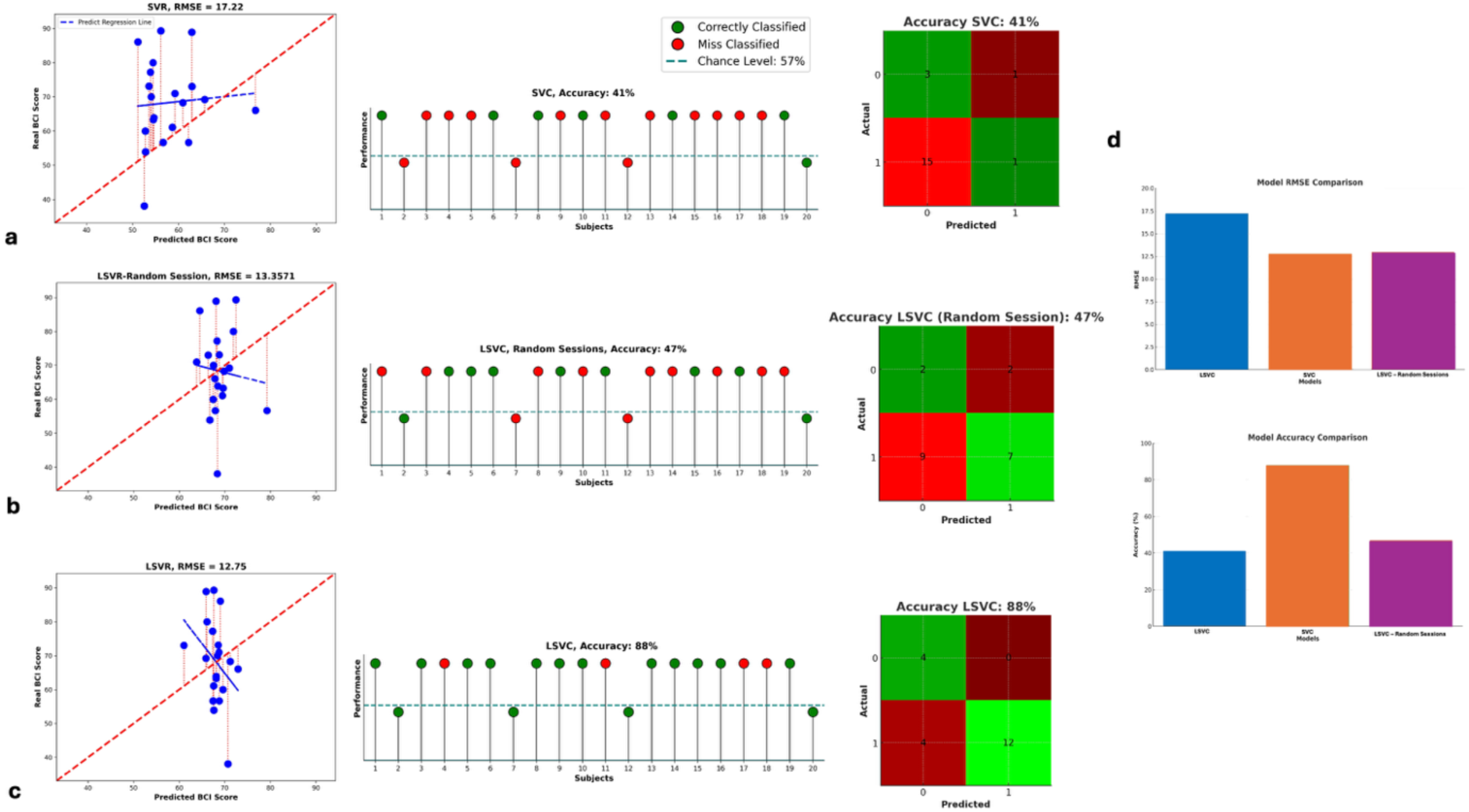
Predictive Model Results. ***a.*** Predictive results using Support Vector Regression (SVR) (left panel) and Support Vector Classification (SVC) (right panel) models. ***b.*** Predictive results using Longitudinal Support Vector Regression (LSVR) (left panel) and Longitudinal Support Vector Classification (LSVC) (right panel) models with sessions in random order. ***c.*** Predictive results using Longitudinal Support Vector Regression (LSVR) (left panel) and Longitudinal Support Vector Classification (LSVC) (right panel) models. **Left Panel: Regression Results:** Each point represents a subject. Red dashed lines indicate the prediction error between actual and predicted values. The bold red line represents the optimal prediction trend, while the blue lines show individual predicted trends. **Right Panel: Classification Results and Confusion Matrix:** Each ball represents a subject. The dashed horizontal line denotes the classification threshold—subjects above this line are predicted to have control. Green balls indicate correct predictions, while red balls represent misclassifications. In the confusion matrix, green cells show correctly classified subjects, and red cells indicate misclassifications. The intensity of each cell’s color corresponds to the number of subjects in that category. ***d.*** Comparison of different models. **Top:** Comparison of the Root Mean Square Error (RMSE) obtained using different models. **Bottom:** Comparison of the accuracy performance obtained using different models. SVC (blue), LSVC (orange), and LSVC with random sessions (purple). All these plots are generated using θ_av_ μ + 3σ and λ_min_av_ : 50ms, and the best-coupled parameters for prediction.

**Supplementary Materials, Figure 2.**
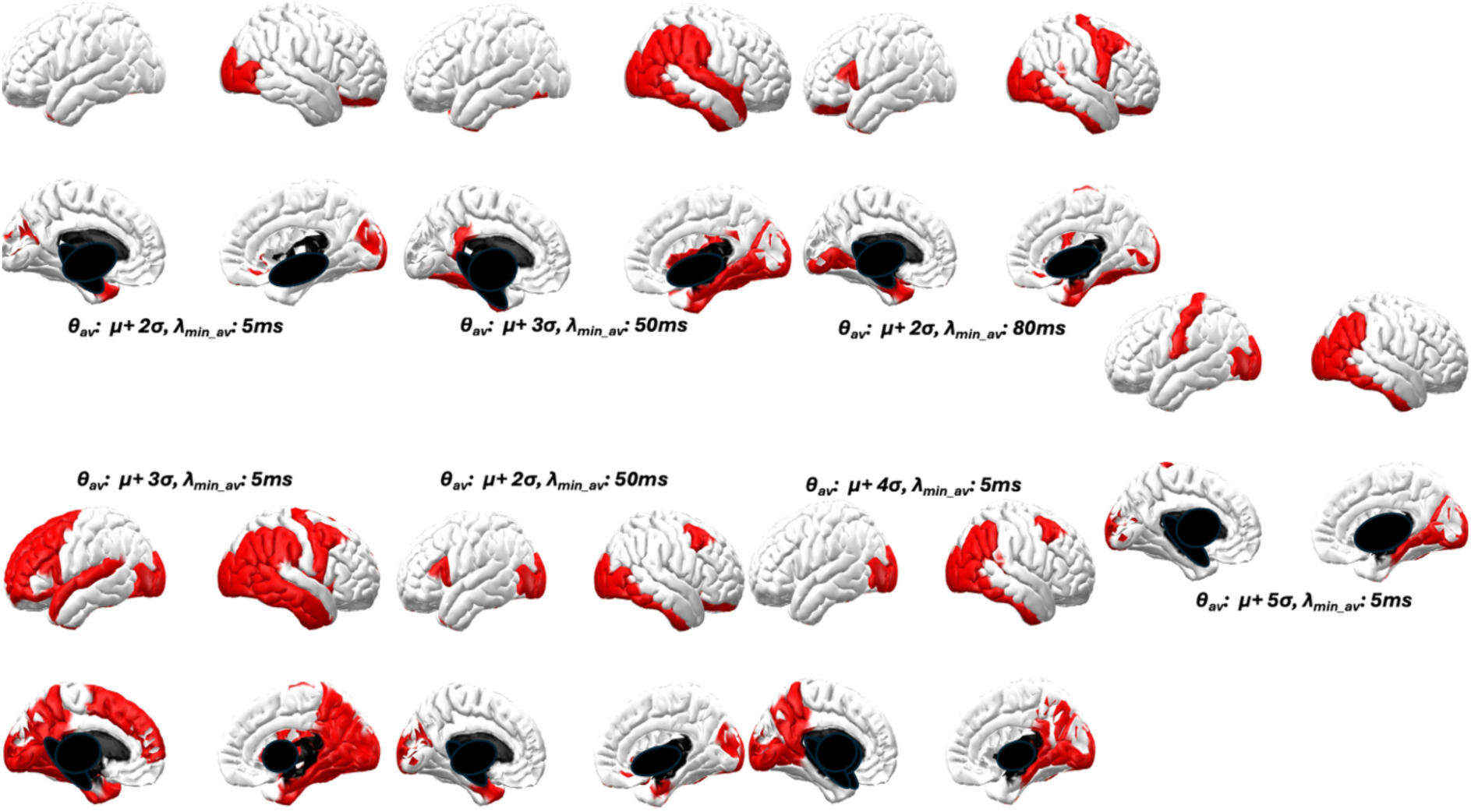
Selected ROIs set for each pair of parameters. In red the ROIs that show significance after ANOVA across all the four sessions of t-values between Motor Imagery and resting condition.

**Supplementary Materials, Table 4.**
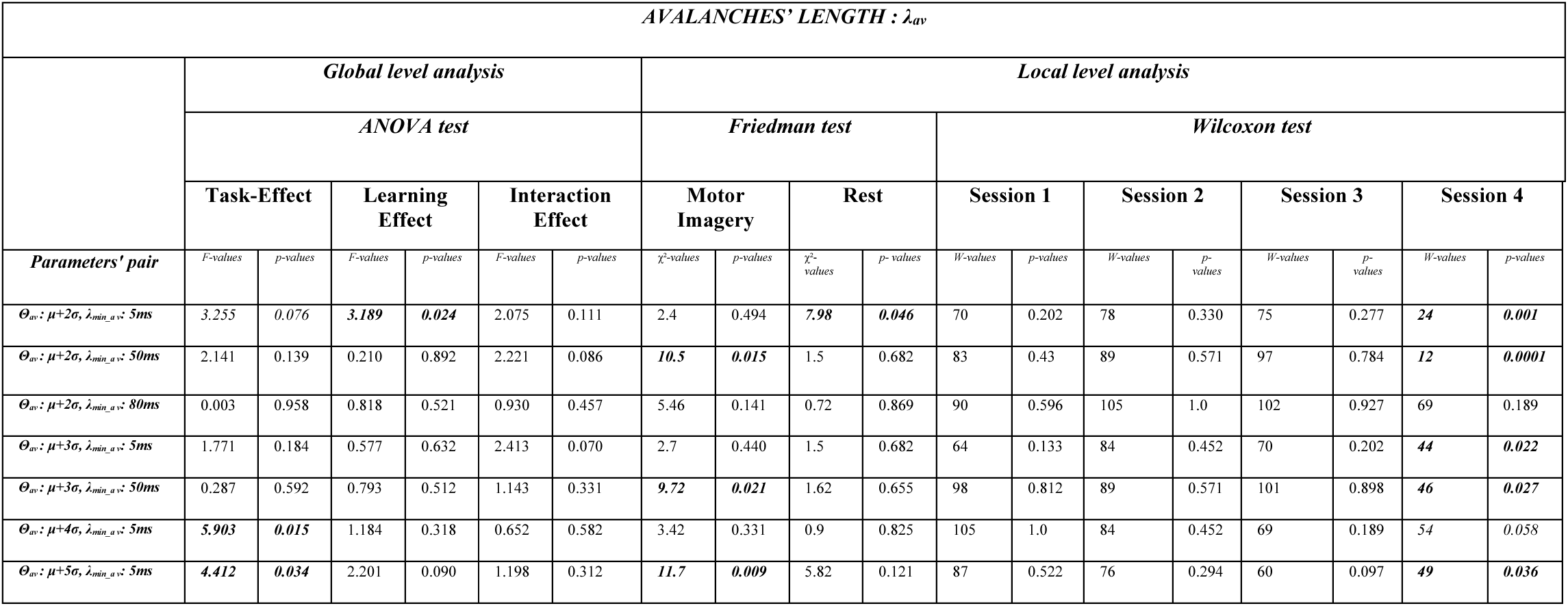
Statistical results of all analysis for Avalanches’ length λ_av_ over all possible parameters’ combinations using a selected set of ROIs. In bold the significant values (p < 0.05)

**Supplementary Materials, Table 5.**
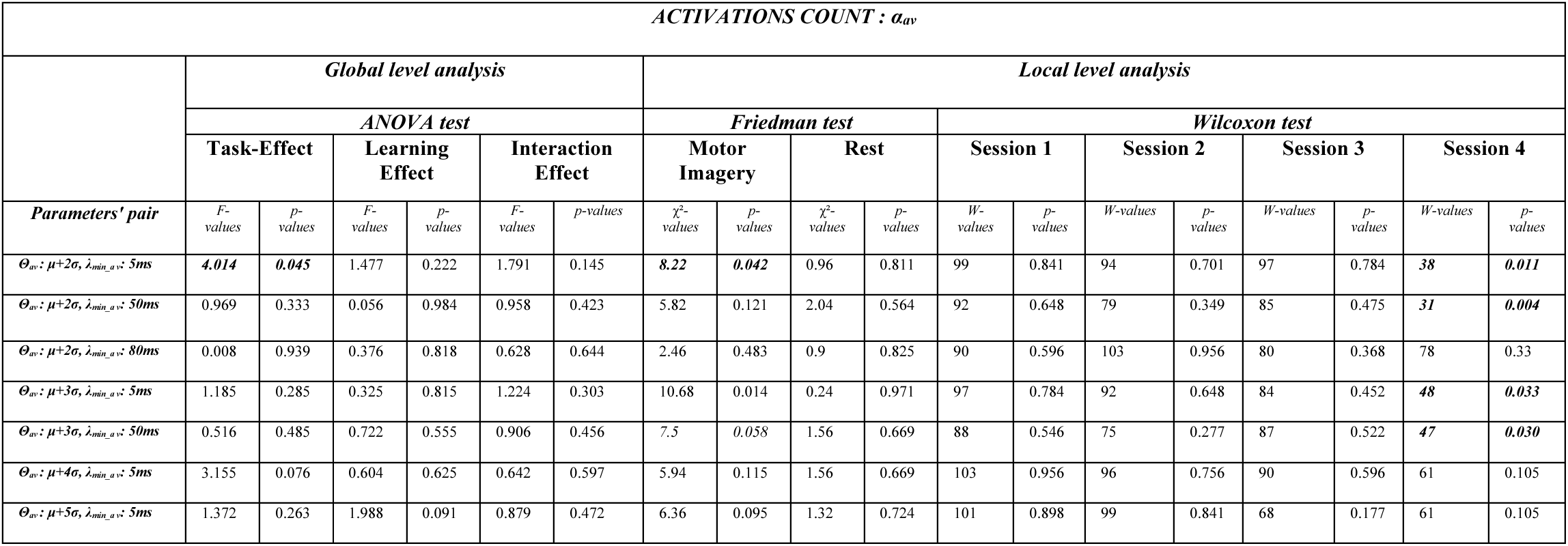
Statistical results of all analysis for Activations count α_av_ over all possible parameters’ combinations using a selected set of ROIs. In bold the significant values (p < 0.05)

**Supplementary Materials, Table 6.**
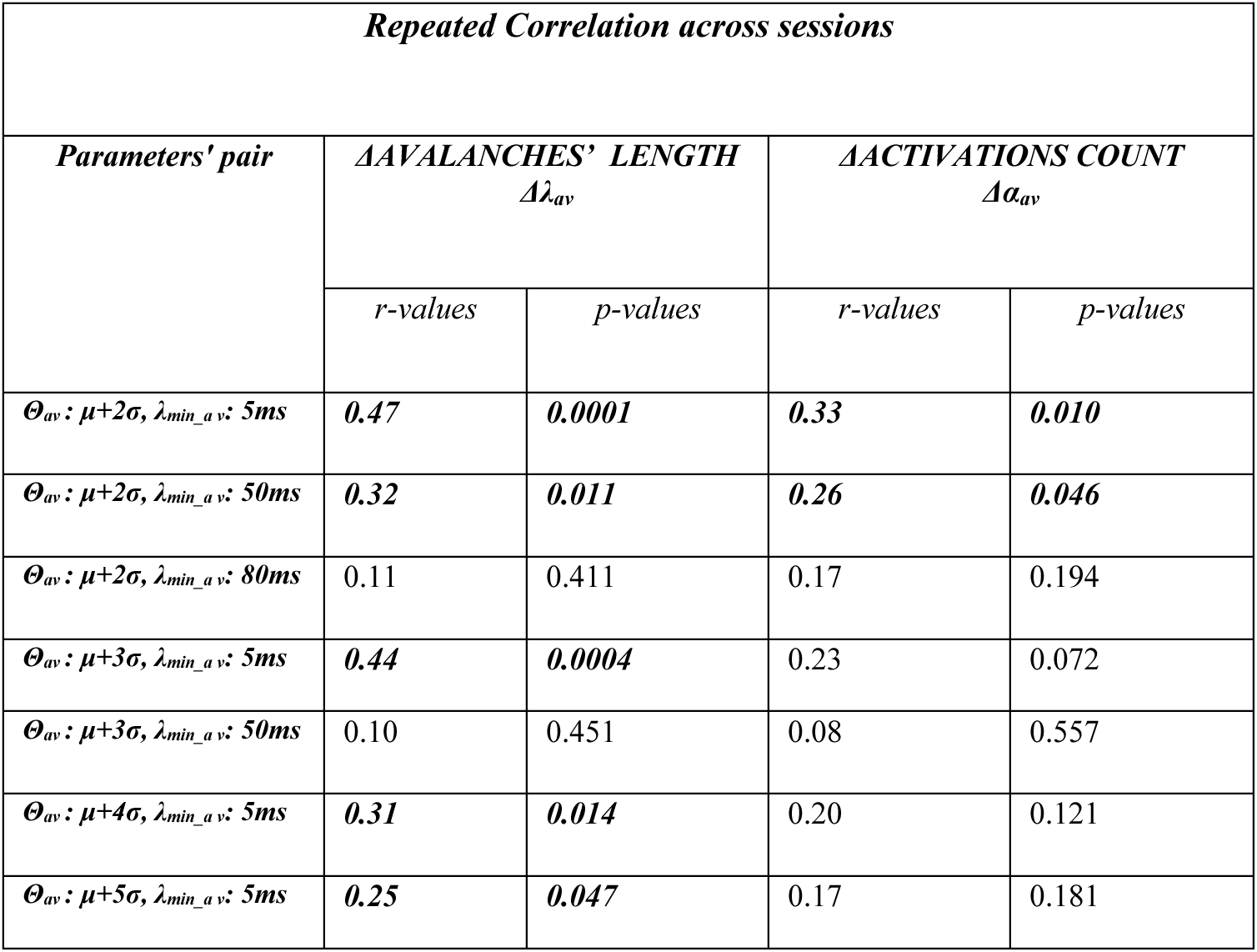
Repeated Correlation results across sessions between both features (Δλ_av_, Δα_av_) and BCI-scores over all possible parameters’ combinations using a selected set of ROIs. *In bold significant correlations (p < 0.05)*

**Supplementary Materials, Figure 3.**
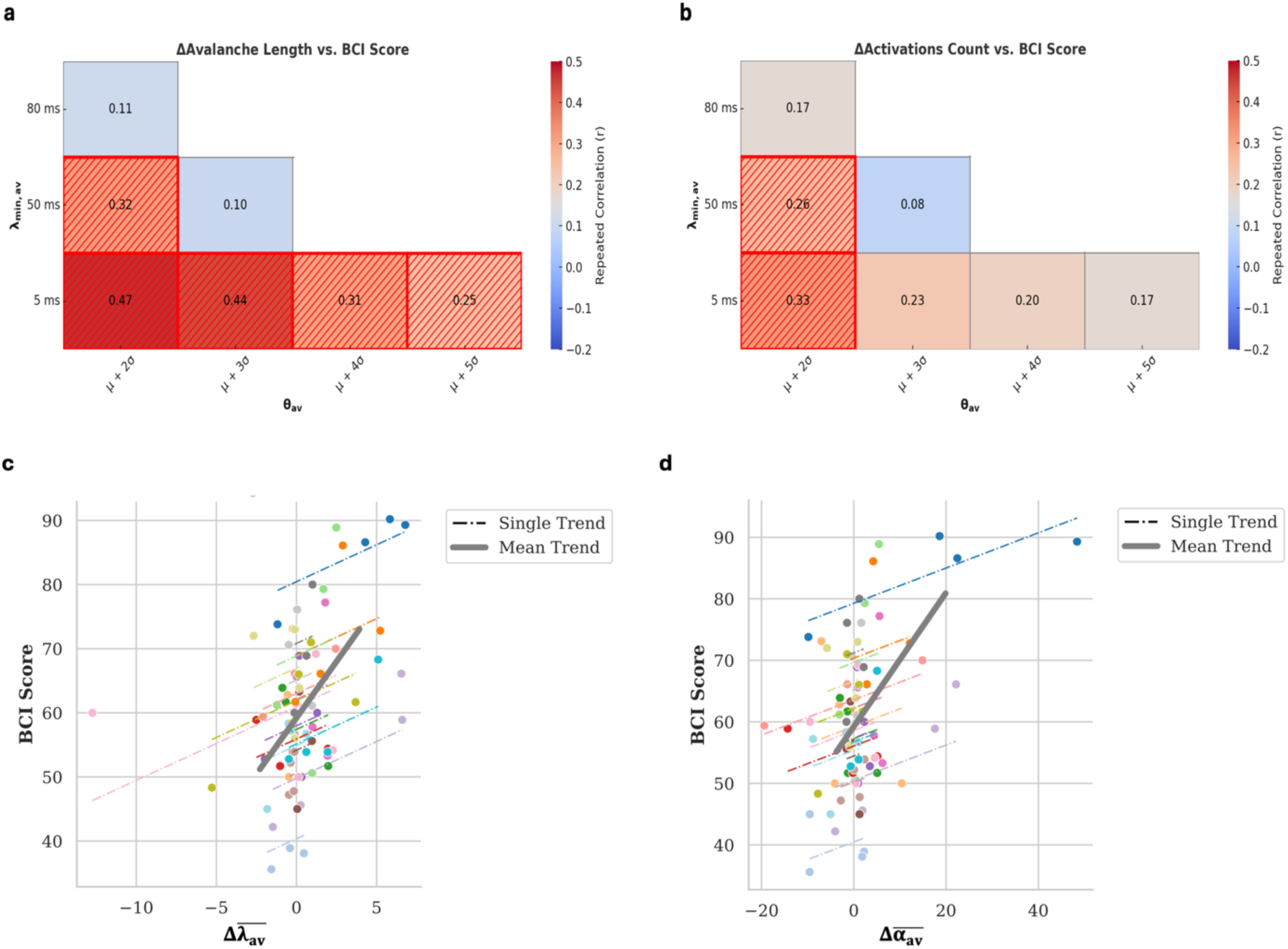
Repeated correlation and trends across different sessions between BCI score and features changes over a selected set of ROIs. *a.* Mean difference of avalanches’ length (Δλ_av_) and *b.* Mean difference activation (Δα_av_) across all tested pairs of parameters (θ_av_, λ_min_av_). Significant correlations (p<0.05) are highlighted using different textures. *c.* Repeated correlations trend across different sessions between BCI-score and Δλ_av_ and *d.* between Δα_av_ and BCI-score. Each coloured dashed line corresponds to one subject while the grey bold line identifies the trend across all the subjects. The pairs of parameters (θ_av_, λ_min_av_) used in *(c)* and *(d)* were those that achieve the best prediction performance: θ_av_: μ+2σ, λ_min_av_ : 50ms.

**Supplementary Materials, Figure 4:**
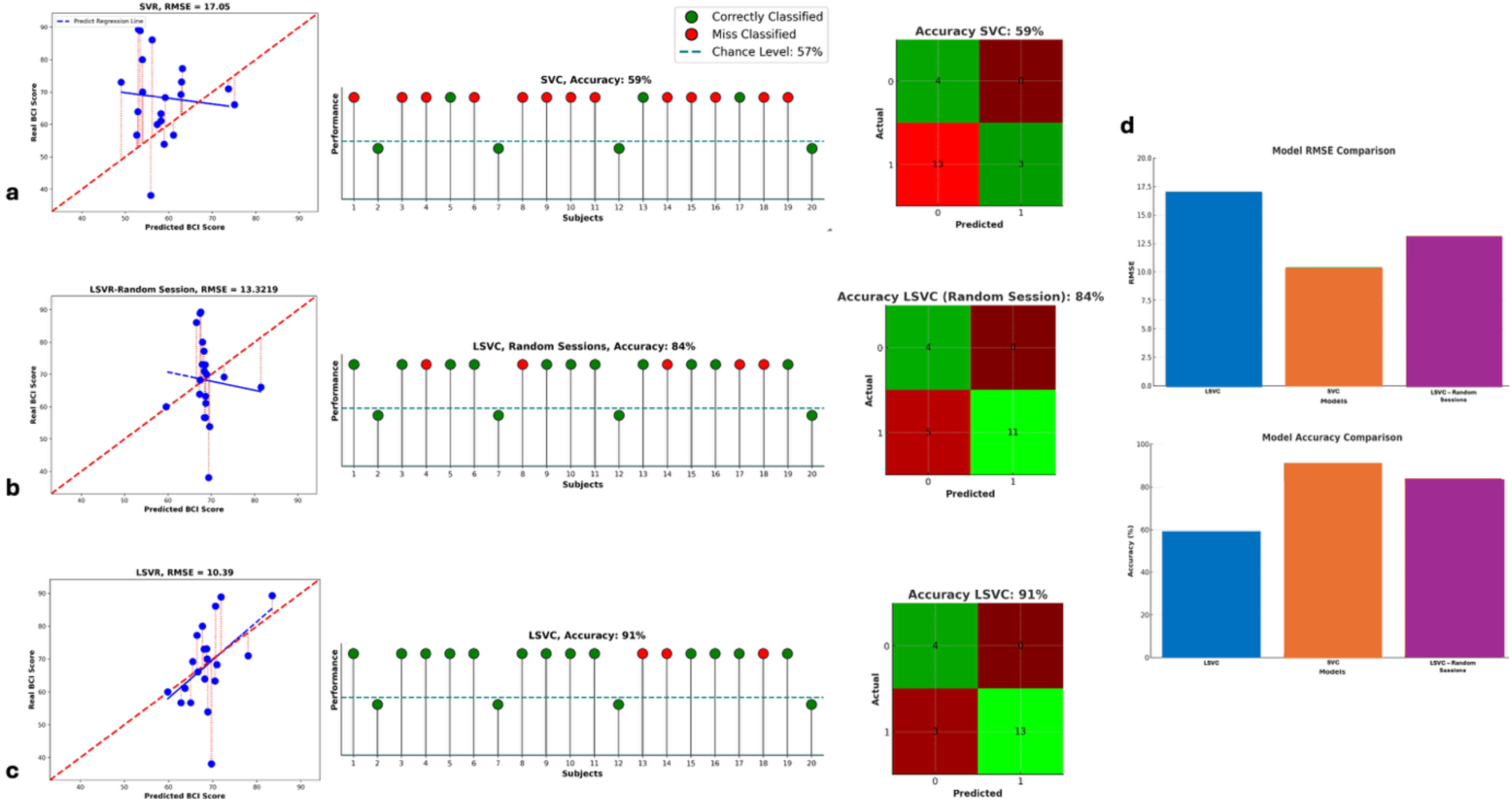
Predictive Model Results over a selected set of ROIs. ***a.*** Predictive results using Support Vector Regression (SVR) (left panel) and Support Vector Classification (SVC) (right panel) models. ***b.*** Predictive results using Longitudinal Support Vector Regression (LSVR) (left panel) and Longitudinal Support Vector Classification (LSVC) (right panel) models with sessions in random order. ***c.*** Predictive results using Longitudinal Support Vector Regression (LSVR) (left panel) and Longitudinal Support Vector Classification (LSVC) (right panel) models. **Left Panel: Regression Results:** Each point represents a subject. Red dashed lines indicate the prediction error between actual and predicted values. The bold red line represents the optimal prediction trend, while the blue lines show individual predicted trends. **Right Panel: Classification Results and Confusion Matrix:** Each ball represents a subject. The dashed horizontal line denotes the classification threshold—subjects above this line are predicted to have control. Green balls indicate correct predictions, while red balls represent misclassifications. In the confusion matrix, green cells show correctly classified subjects, and red cells indicate misclassifications. The intensity of each cell’s color corresponds to the number of subjects in that category. ***d.*** Comparison of different models. **Top:** Comparison of the Root Mean Square Error (RMSE) obtained using different models. **Bottom:** Comparison of the accuracy performance obtained using different models. SVC (blue), LSVC (orange), and LSVC with random sessions (purple). All these plots are generated using θ_av_ μ + 2σ and λ_min_av_ : 50ms, and the best-coupled parameters for prediction.

**Supplementary Materials, Figure 5.**
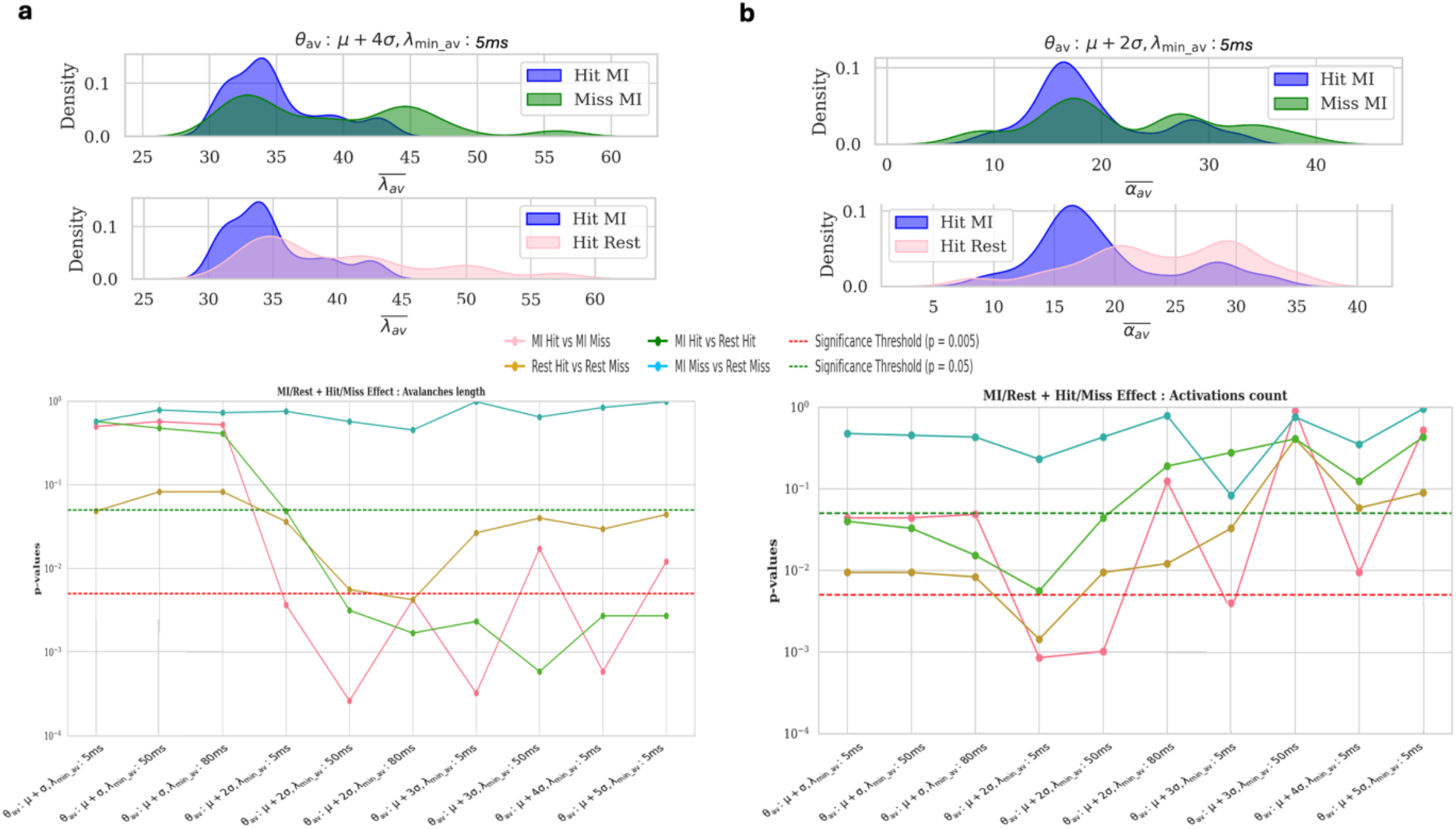
Analysis of Mean Avalanche Length and Activations in Hit vs. Miss Trials. ***a.* Mean Avalanche Length in Hit Trials vs. Miss Trials**: **Top:** Probability density functions representing: i) Hit Motor Imagery (MI) trials (in blue) vs. Hit Rest trials (in pink) on the left; ii) Hit MI trials (in blue) vs. Miss MI trials (in green) on the right. **Bottom:** Trends of p-values from pairwise comparisons of mean avalanche length across brain states (MI vs. Rest) and conditions (Hit vs. Miss) across possible coupled parameters. Dashed horizontal lines indicate significance thresholds (green: p < 0.05: red: p < 0.001, Bonferroni-corrected). ***b.* Activations in Hit Trials vs. Miss Trials:** **Top:** Probability density functions representing: i) Hit Motor Imagery (MI) trials (in blue) vs. Hit Rest trials (in pink) on the left; ii) Hit MI trials (in blue) vs. Miss MI trials (in green) on the right. **Bottom:** Trends of p-values from pairwise comparisons of activation counts across brain states (MI vs. Rest) and conditions (Hit vs. Miss) across possible coupled parameters. Dashed horizontal lines indicate significance thresholds (green:p < 0.05, red: p < 0.001, Bonferroni-corrected). All these analyses are performed only on the last training session.

**Supplementary Materials, Figure 6.**
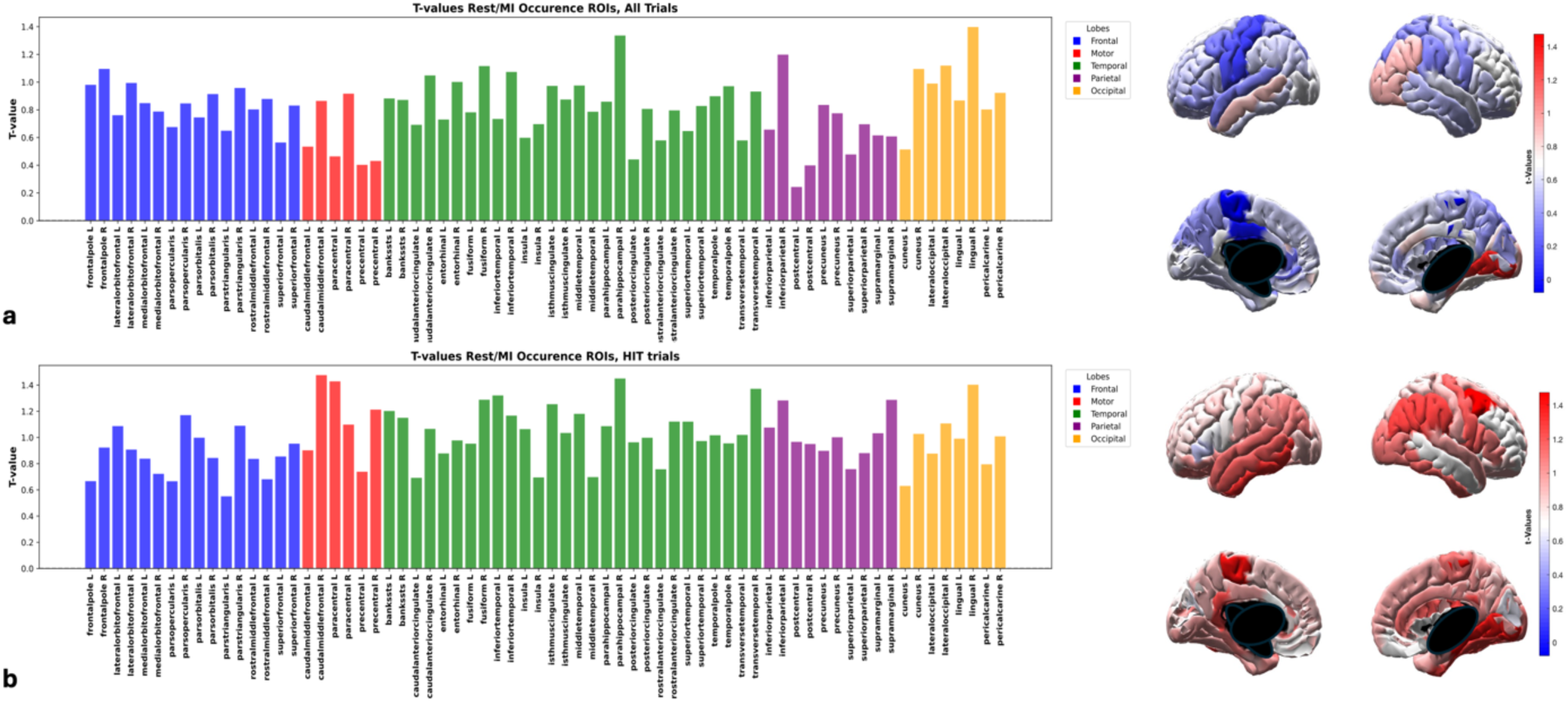
t-values Comparing Rest and Motor Imagery (MI) Conditions. ***a.*** t-values calculated for the occurrence of each specific Region of Interest (ROI) across different trials, considering all trials. ***b.*** t-values calculated for the occurrence of each specific ROI across different trials, considering only those trials in which the subject successfully controlled the device. In the brain plots, the color map reflects the magnitude of the t-values, while in the bar plots, it corresponds to the height of each bar. Brain regions are color-coded as follows: red for the motor cortex, green for the temporal lobe, yellow for the occipital lobe, blue for the frontal lobe, purple for the parietal lobe. All these analyses were performed only on the last training session and one specific parameter combination: θ_av_: μ + 4σ, λ_min_av_: 2.

