## Supplementary Materials for "Neuronal avalanches as a predictive biomarker for guiding tailored BCI training programs"

| *AVALANCHES' LENGTH : $\lambda_{av}$* | | | | | | | | | | | | | | | | | | |
|---|---|---|---|---|---|---|---|---|---|---|---|---|---|---|---|---|---|---|
| | ***Global level analysis*** | | | | | | ***Local level analysis*** | | | | | | | | | | | |
| | ***ANOVA test*** | | | | | | ***Friedman test*** | | | | ***Wilcoxon test*** | | | | | | | |
| | **Task-Effect** | | **Learning Effect** | | **Interaction Effect** | | **Motor Imagery** | | **Rest** | | **Session 1** | | **Session 2** | | **Session 3** | | **Session 4** | |
| ***Parameters' pair*** | *F-values* | *p-values* | *F-values* | *p-values* | *F-values* | *p-values* | *$\chi^2$-values* | *p-values* | *$\chi^2$-values* | *p- values* | *W-values* | *p-values* | *W-values* | *p-values* | *W-values* | *p-values* | *W-values* | *p-values* |
| ***$\Theta_{av}$ : $\mu+\sigma$, $\lambda_{min\_av}$: 5ms*** | 1.450 | 0.243 | 1.208 | 0.3203 | 0.740 | 0.540 | 2.82 | 0.4202 | 2.82 | 0.420 | 71.0 | 0.216 | 103.0 | 0.956 | 56.0 | 0.070 | 68.0 | 0.177 |
| ***$\Theta_{av}$ : $\mu+\sigma$, $\lambda_{min\_av}$: 50ms*** | 1.308 | 0.259 | 1.192 | 0.314 | 0.752 | 0.535 | 2.82 | 0.420 | 1.98 | 0.577 | 69.0 | 0.189 | 81.0 | 0.388 | 59.0 | 0.090 | 76.0 | 0.294 |
| ***$\Theta_{av}$ : $\mu+\sigma$, $\lambda_{min\_av}$: 80ms*** | 1.216 | 0.275 | 1.190 | 0.323 | 0.766 | 0.516 | 5.34 | 0.149 | 3.06 | 0.390 | 68.0 | 0.177 | 65.0 | 0.143 | 61.0 | 0.106 | 80.0 | 0.368 |
| ***$\Theta_{av}$ : $\mu+2\sigma$, $\lambda_{min\_av}$: 5ms*** | 1.378 | 0.252 | 0.857 | 0.486 | 0.414 | 0.753 | 4.98 | 0.173 | 1.38 | 0.710 | 102.0 | 0.927 | 71.0 | 0.216 | 92.0 | 0.648 | 96.0 | 0.756 |
| ***$\Theta_{av}$ : $\mu+2\sigma$, $\lambda_{min\_av}$: 50ms*** | 0.179 | 0.680 | 0.850 | 0.482 | 0.397 | 0.770 | 1.68 | 0.641 | 1.02 | 0.796 | 78.0 | 0.330 | 83.0 | 0.430 | 78.0 | 0.330 | 79.0 | 0.349 |
| ***$\Theta_{av}$ : $\mu+2\sigma$, $\lambda_{min\_av}$: 80ms*** | 0.037 | 0.849 | 0.458 | 0.726 | 0.573 | 0.645 | 6.66 | 0.084 | 2.34 | 0.505 | 61.0 | 0.105 | 105.0 | 1.0 | 69.0 | 0.189 | 76.0 | 0.294 |
| ***$\Theta_{av}$ : $\mu+3\sigma$, $\lambda_{min\_av}$: 5ms*** | 0.021 | 0.886 | 0.288 | 0.840 | 1.990 | 0.121 | 4.5 | 0.212 | 7.38 | 0.061 | 77.0 | 0.312 | 85.0 | 0.475 | 73.0 | 0.246 | 53.0 | 0.053 |
| ***$\Theta_{av}$ : $\mu+3\sigma$, $\lambda_{min\_av}$: 50ms*** | 0.005 | 0.943 | 0.670 | 0.575 | 2.182 | 0.093 | ***9.66*** | ***0.022*** | 0.42 | 0.936 | 57.0 | 0.076 | 75.0 | 0.277 | 78.0 | 0.330 | ***34.0*** | ***0.006*** |
| ***$\Theta_{av}$ : $\mu+4\sigma$, $\lambda_{min\_av}$: 5ms*** | 3.406 | 0.067 | 0.628 | 0.597 | 1.523 | 0.205 | 5.1 | 0.165 | 1.2 | 0.753 | 76.0 | 0.294 | 88.0 | 0.546 | 75.0 | 0.277 | 56.0 | 0.070 |
| ***$\Theta_{av}$ : $\mu+5\sigma$, $\lambda_{min\_av}$: 5ms*** | 3.560 | 0.064 | 0.965 | 0.422 | 0.933 | 0.429 | ***8.7*** | ***0.034*** | 0.78 | 0.854 | 97.0 | 0.784 | 89.0 | 0.571 | 88.0 | 0.546 | ***52.0*** | ***0.048*** |

**Supplementary Materials, Table 1. Statistical results of all analysis for Avalanches' length $\lambda_{av}$ over all possible parameters' combinations.** In bold the significant values ($p < 0.05$)

| *ACTIVATIONS COUNT : $\alpha_{av}$* | | | | | | | | | | | | | | | | | | |
|---|---|---|---|---|---|---|---|---|---|---|---|---|---|---|---|---|---|---|
| | ***Global level analysis*** | | | | | | ***Local level analysis*** | | | | | | | | | | | |
| | ***ANOVA test*** | | | | | | ***Friedman test*** | | | | ***Wilcoxon test*** | | | | | | | |
| | **Task-Effect** | | **Learning Effect** | | **Interaction Effect** | | **Motor Imagery** | | **Rest** | | **Session 1** | | **Session 2** | | **Session 3** | | **Session 4** | |
| ***Parameters' pair*** | *F-values* | *p-values* | *F-values* | *p-values* | *F-values* | *p-values* | *$\chi^2$ values* | *p-values* | *$\chi^2$-values* | *p- values* | *W-values* | *p-values* | *W-values* | *p-values* | *W-values* | *p-values* | *W-values* | *p-values* |
| ***$\Theta_{av}$ : $\mu+\sigma$, $\lambda_{min\_av}$: 5ms*** | 0.964 | 0.329 | 1.371 | 0.253 | 0.667 | 0.588 | 4.38 | 0.223 | 4.02 | 0.259 | 80 | 0.368 | 81 | 0.388 | 93 | 0.674 | 79 | 0.349 |
| ***$\Theta_{av}$ : $\mu+\sigma$, $\lambda_{min\_av}$: 50ms*** | 0.904 | 0.348 | 1.468 | 0.229 | 0.676 | 0.576 | 5.28 | 0.152 | 5.46 | 0.141 | 79 | 0.349 | 79 | 0.349 | 96 | 0.756 | 82 | 0.409 |
| ***$\Theta_{av}$ : $\mu+\sigma$, $\lambda_{min\_av}$: 80ms*** | 0.856 | 0.379 | 1.522 | 0.215 | 0.683 | 0.574 | 4.5 | 0.212 | 3.96 | 0.266 | 80 | 0.368 | 81 | 0.388 | 99 | 0.841 | 80 | 0.368 |
| ***$\Theta_{av}$ : $\mu+2\sigma$, $\lambda_{min\_av}$: 5ms*** | 0.056 | 0.812 | 0.841 | 0.482 | 0.535 | 0.672 | 5.76 | 0.124 | 1.86 | 0.602 | 84 | 0.452 | 99 | 0.841 | 72 | 0.231 | 79 | 0.349 |
| ***$\Theta_{av}$ : $\mu+2\sigma$, $\lambda_{min\_av}$: 50ms*** | 0.285 | 0.597 | 0.829 | 0.489 | 0.899 | 0.451 | 3.12 | 0.374 | 1.62 | 0.655 | 82 | 0.409 | 92 | 0.648 | 85 | 0.475 | 75 | 0.277 |
| ***$\Theta_{av}$ : $\mu+2\sigma$, $\lambda_{min\_av}$: 80ms*** | 0.609 | 0.436 | 0.597 | 0.627 | 1.106 | 0.344 | 4.2 | 0.241 | 1.86 | 0.602 | 80 | 0.368 | 93 | 0.674 | 84 | 0.452 | 69 | 0.189 |
| ***$\Theta_{av}$ : $\mu+3\sigma$, $\lambda_{min\_av}$: 5ms*** | 0.426 | 0.517 | 0.323 | 0.818 | 1.077 | 0.367 | ***12.78*** | ***0.005*** | 0.78 | 0.854 | 105 | 1.0 | 92 | 0.648 | 86 | 0.498 | ***47*** | ***0.030*** |
| ***$\Theta_{av}$ : $\mu+3\sigma$, $\lambda_{min\_av}$: 50ms*** | 0.15 | 0.699 | 0.218 | 0.887 | 1.2 | 0.306 | 5.58 | 0.134 | 0.96 | 0.811 | 94 | 0.701 | 84 | 0.452 | 93 | 0.674 | ***52*** | ***0.048*** |
| ***$\Theta_{av}$ : $\mu+4\sigma$, $\lambda_{min\_av}$: 5ms*** | 0.357 | 0.561 | 0.643 | 0.612 | 0.552 | 0.665 | ***9.42*** | ***0.024*** | 0.18 | 0.981 | 94 | 0.701 | 87 | 0.522 | 69 | 0.189 | 73 | 0.246 |
| ***$\Theta_{av}$ : $\mu+5\sigma$, $\lambda_{min\_av}$: 5ms*** | 0.079 | 0.794 | 1.018 | 0.39 | 0.355 | 0.797 | 3.0 | 0.392 | 0.3 | 0.96 | 98 | 0.812 | 83 | 0.43 | 68 | 0.177 | 76 | 0.294 |

**Supplementary Materials, Table 2. Statistical results of all analysis for Activations count $\alpha_{av}$ over all possible parameters' combinations.** In bold the significant values ($p < 0.05$)

| ***Repeated Correlation across sessions*** | | | | |
|---|---|---|---|---|
| ***Parameters' pair*** | ***ΔAVALANCHES' LENGTH*** ***$\Delta\lambda_{av}$*** | | ***ΔACTIVATIONS COUNT*** ***$\Delta\alpha_{av}$*** | |
| | *r-values* | *p-values* | *r-values* | *p-values* |
| ***$\Theta_{av}$ : $\mu+\sigma$, $\lambda_{min\_av}$: 5ms*** | -0.01 | 0.905 | 0.004 | 0.976 |
| ***$\Theta_{av}$ : $\mu+\sigma$, $\lambda_{min\_av}$: 50ms*** | -0.01 | 0.917 | 0.01 | 0.937 |
| ***$\Theta_{av}$ : $\mu+\sigma$, $\lambda_{min\_av}$: 80ms*** | -0.02 | 0.943 | 0.01 | 0.917 |
| ***$\Theta_{av}$ : $\mu+2\sigma$, $\lambda_{min\_av}$: 5ms*** | -0.12 | 0.343 | 0.20 | 0.120 |
| ***$\Theta_{av}$ : $\mu+2\sigma$, $\lambda_{min\_av}$: 50ms*** | 0.02 | 0.877 | ***0.29*** | ***0.022*** |
| ***$\Theta_{av}$ : $\mu+2\sigma$, $\lambda_{min\_av}$: 80ms*** | 0.09 | 0.484 | ***0.26*** | ***0.040*** |
| ***$\Theta_{av}$ : $\mu+3\sigma$, $\lambda_{min\_av}$: 5ms*** | ***0.27*** | ***0.033*** | ***0.28*** | ***0.027*** |
| ***$\Theta_{av}$ : $\mu+3\sigma$, $\lambda_{min\_av}$: 50ms*** | ***0.39*** | ***0.002*** | ***0.28*** | ***0.028*** |
| ***$\Theta_{av}$ : $\mu+4\sigma$, $\lambda_{min\_av}$: 5ms*** | ***0.48*** | ***0.0001*** | 0.20 | 0.122 |
| ***$\Theta_{av}$ : $\mu+5\sigma$, $\lambda_{min\_av}$: 5ms*** | ***0.47*** | ***0.0002*** | 0.15 | 0.247 |

**Supplementary Materials, Table 3. Repeated Correlation results across sessions between both features ($\Delta\lambda_{av}$ , $\Delta\alpha_{av}$) and BCI-scores over all possible parameters' combinations.** In bold significant correlations ($p < 0.05$)

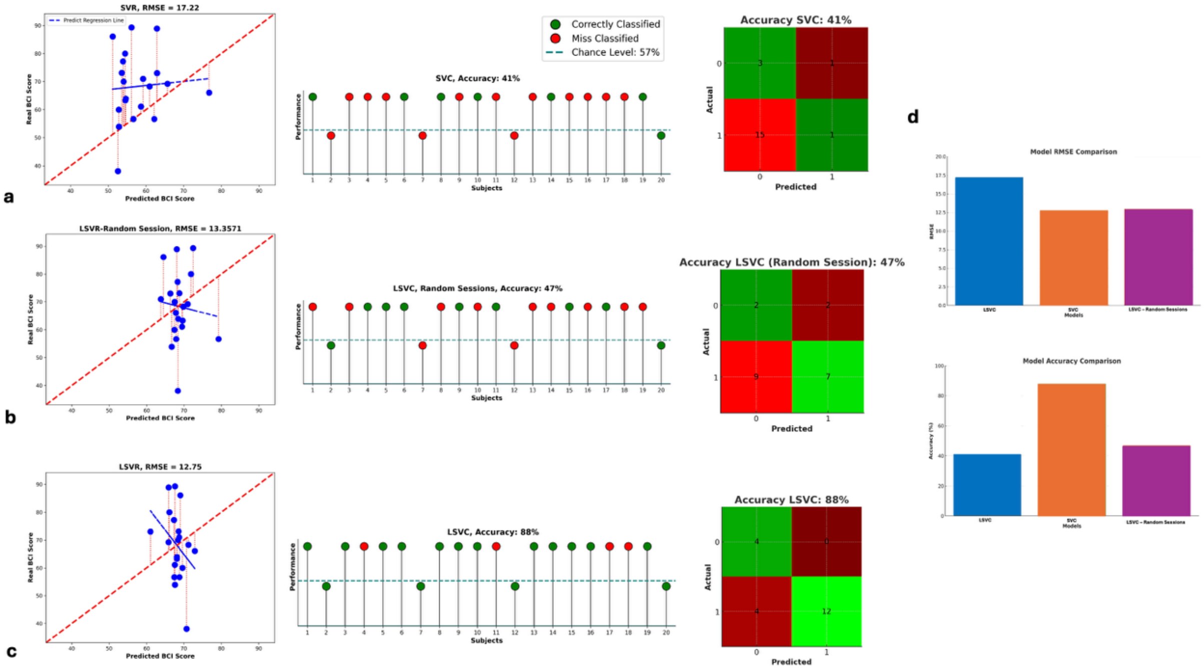

**Supplementary Materials, Figure 1. Predictive Model Results**.

***a.*** Predictive results using Support Vector Regression (SVR) (left panel) and Support Vector Classification (SVC) (right panel) models.

***b.*** Predictive results using Longitudinal Support Vector Regression (LSVR) (left panel) and Longitudinal Support Vector Classification (LSVC) (right panel) models with sessions in random order.

***c.*** Predictive results using Longitudinal Support Vector Regression (LSVR) (left panel) and Longitudinal Support Vector Classification (LSVC) (right panel) models.

> **Left Panel:** Regression Results: Each point represents a subject. Red dashed lines indicate the prediction error between actual and predicted values. The bold red line represents the optimal prediction trend, while the blue lines show individual predicted trends.
>
> **Right Panel:** Classification Results and Confusion Matrix**:** Each ball represents a subject. The dashed horizontal line denotes the classification threshold—subjects above this line are predicted to have control. Green balls indicate correct predictions, while red balls represent misclassifications. In the confusion matrix, green cells show correctly classified subjects, and red cells indicate misclassifications. The intensity of each cell's color corresponds to the number of subjects in that category.

***d.*** Comparison of different models.

> **Top:** Comparison of the Root Mean Square Error (RMSE) obtained using different models.
>
> **Bottom:** Comparison of the accuracy performance obtained using different models. SVC (blue), LSVC (orange), and LSVC with random sessions (purple).

All these plots are generated using $\theta_{av}$ $\mu + 3\sigma$ and $\lambda_{min\_av}$ : 50ms, and the best-coupled parameters for prediction.

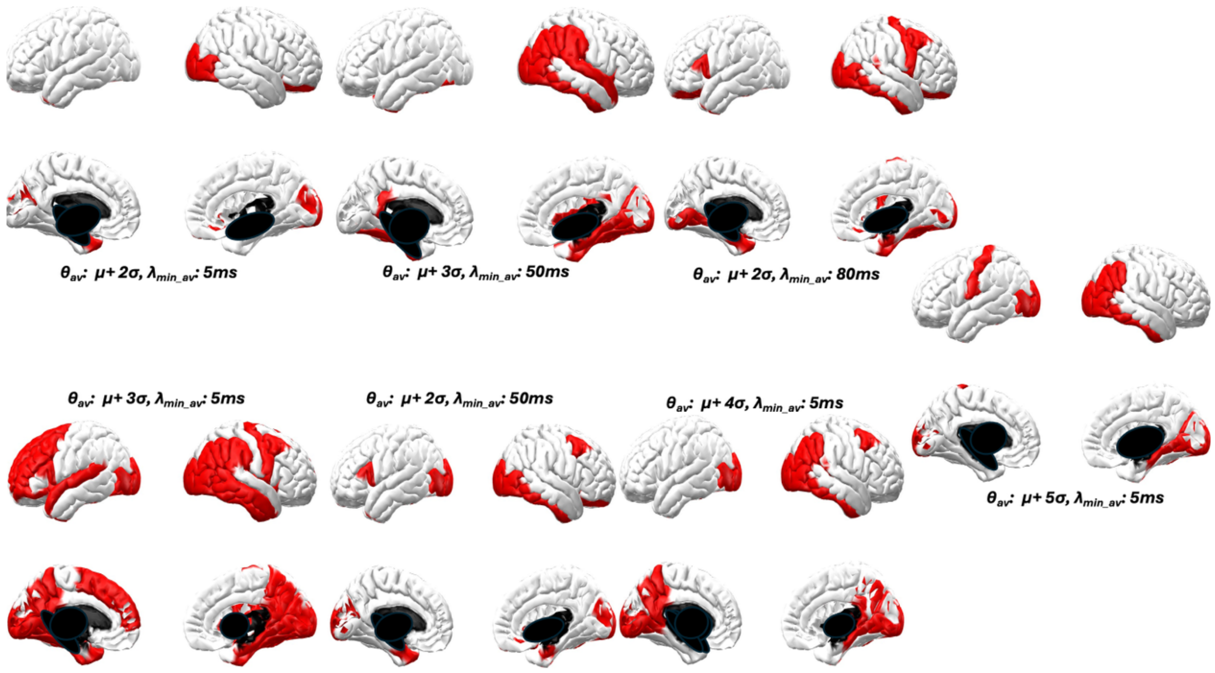

**Supplementary Materials, Figure 2. Selected ROIs set for each pair of parameters.** In red the ROIs that show significance after ANOVA across all the four sessions of t-values between Motor Imagery and resting condition.

| *AVALANCHES' LENGTH : $\lambda_{av}$* | | | | | | | | | | | | | | | | | | |
|---|---|---|---|---|---|---|---|---|---|---|---|---|---|---|---|---|---|---|
| | *Global level analysis* | | | | | | *Local level analysis* | | | | | | | | | | | |
| | *ANOVA test* | | | | | | *Friedman test* | | | | *Wilcoxon test* | | | | | | | |
| | **Task-Effect** | | **Learning Effect** | | **Interaction Effect** | | **Motor Imagery** | | **Rest** | | **Session 1** | | **Session 2** | | **Session 3** | | **Session 4** | |
| ***Parameters' pair*** | *F-values* | *p-values* | *F-values* | *p-values* | *F-values* | *p-values* | *$\chi^2$-values* | *p-values* | *$\chi^2$-values* | *p- values* | *W-values* | *p-values* | *W-values* | *p-values* | *W-values* | *p-values* | *W-values* | *p-values* |
| ***$\Theta_{av}$ : $\mu+2\sigma$, $\lambda_{min\_av}$: 5ms*** | *3.255* | *0.076* | ***3.189*** | ***0.024*** | 2.075 | 0.111 | 2.4 | 0.494 | ***7.98*** | ***0.046*** | 70 | 0.202 | 78 | 0.330 | 75 | 0.277 | ***24*** | ***0.001*** |
| ***$\Theta_{av}$ : $\mu+2\sigma$, $\lambda_{min\_av}$: 50ms*** | 2.141 | 0.139 | 0.210 | 0.892 | 2.221 | 0.086 | ***10.5*** | ***0.015*** | 1.5 | 0.682 | 83 | 0.43 | 89 | 0.571 | 97 | 0.784 | ***12*** | ***0.0001*** |
| ***$\Theta_{av}$ : $\mu+2\sigma$, $\lambda_{min\_av}$: 80ms*** | 0.003 | 0.958 | 0.818 | 0.521 | 0.930 | 0.457 | 5.46 | 0.141 | 0.72 | 0.869 | 90 | 0.596 | 105 | 1.0 | 102 | 0.927 | 69 | 0.189 |
| ***$\Theta_{av}$ : $\mu+3\sigma$, $\lambda_{min\_av}$: 5ms*** | 1.771 | 0.184 | 0.577 | 0.632 | 2.413 | 0.070 | 2.7 | 0.440 | 1.5 | 0.682 | 64 | 0.133 | 84 | 0.452 | 70 | 0.202 | ***44*** | ***0.022*** |
| ***$\Theta_{av}$ : $\mu+3\sigma$, $\lambda_{min\_av}$: 50ms*** | 0.287 | 0.592 | 0.793 | 0.512 | 1.143 | 0.331 | ***9.72*** | ***0.021*** | 1.62 | 0.655 | 98 | 0.812 | 89 | 0.571 | 101 | 0.898 | ***46*** | ***0.027*** |
| ***$\Theta_{av}$ : $\mu+4\sigma$, $\lambda_{min\_av}$: 5ms*** | ***5.903*** | ***0.015*** | 1.184 | 0.318 | 0.652 | 0.582 | 3.42 | 0.331 | 0.9 | 0.825 | 105 | 1.0 | 84 | 0.452 | 69 | 0.189 | *54* | *0.058* |
| ***$\Theta_{av}$ : $\mu+5\sigma$, $\lambda_{min\_av}$: 5ms*** | ***4.412*** | ***0.034*** | 2.201 | 0.090 | 1.198 | 0.312 | ***11.7*** | ***0.009*** | 5.82 | 0.121 | 87 | 0.522 | 76 | 0.294 | 60 | 0.097 | ***49*** | ***0.036*** |

**Supplementary Materials, Table 4. Statistical results of all analysis for Avalanches' length $\lambda_{av}$ over all possible parameters' combinations using a selected set of ROIs.** In bold the significant values ($p < 0.05$)

| *ACTIVATIONS COUNT : $\alpha_{av}$* | | | | | | | | | | | | | | | | | | |
|---|---|---|---|---|---|---|---|---|---|---|---|---|---|---|---|---|---|---|
| | *Global level analysis* | | | | | | *Local level analysis* | | | | | | | | | | | |
| | *ANOVA test* | | | | | | *Friedman test* | | | | *Wilcoxon test* | | | | | | | |
| | **Task-Effect** | | **Learning Effect** | | **Interaction Effect** | | **Motor Imagery** | | **Rest** | | **Session 1** | | **Session 2** | | **Session 3** | | **Session 4** | |
| ***Parameters' pair*** | *F-values* | *p-values* | *F-values* | *p-values* | *F-values* | *p-values* | *$\chi^2$-values* | *p-values* | *$\chi^2$-values* | *p-values* | *W-values* | *p-values* | *W-values* | *p-values* | *W-values* | *p-values* | *W-values* | *p-values* |
| ***$\Theta_{av}$ : $\mu+2\sigma$, $\lambda_{min\_av}$: 5ms*** | ***4.014*** | ***0.045*** | 1.477 | 0.222 | 1.791 | 0.145 | ***8.22*** | ***0.042*** | 0.96 | 0.811 | 99 | 0.841 | 94 | 0.701 | 97 | 0.784 | ***38*** | ***0.011*** |
| ***$\Theta_{av}$ : $\mu+2\sigma$, $\lambda_{min\_av}$: 50ms*** | 0.969 | 0.333 | 0.056 | 0.984 | 0.958 | 0.423 | 5.82 | 0.121 | 2.04 | 0.564 | 92 | 0.648 | 79 | 0.349 | 85 | 0.475 | ***31*** | ***0.004*** |
| ***$\Theta_{av}$ : $\mu+2\sigma$, $\lambda_{min\_av}$: 80ms*** | 0.008 | 0.939 | 0.376 | 0.818 | 0.628 | 0.644 | 2.46 | 0.483 | 0.9 | 0.825 | 90 | 0.596 | 103 | 0.956 | 80 | 0.368 | 78 | 0.33 |
| ***$\Theta_{av}$ : $\mu+3\sigma$, $\lambda_{min\_av}$: 5ms*** | 1.185 | 0.285 | 0.325 | 0.815 | 1.224 | 0.303 | 10.68 | 0.014 | 0.24 | 0.971 | 97 | 0.784 | 92 | 0.648 | 84 | 0.452 | ***48*** | ***0.033*** |
| ***$\Theta_{av}$ : $\mu+3\sigma$, $\lambda_{min\_av}$: 50ms*** | 0.516 | 0.485 | 0.722 | 0.555 | 0.906 | 0.456 | *7.5* | *0.058* | 1.56 | 0.669 | 88 | 0.546 | 75 | 0.277 | 87 | 0.522 | ***47*** | ***0.030*** |
| ***$\Theta_{av}$ : $\mu+4\sigma$, $\lambda_{min\_av}$: 5ms*** | 3.155 | 0.076 | 0.604 | 0.625 | 0.642 | 0.597 | 5.94 | 0.115 | 1.56 | 0.669 | 103 | 0.956 | 96 | 0.756 | 90 | 0.596 | 61 | 0.105 |
| ***$\Theta_{av}$ : $\mu+5\sigma$, $\lambda_{min\_av}$: 5ms*** | 1.372 | 0.263 | 1.988 | 0.091 | 0.879 | 0.472 | 6.36 | 0.095 | 1.32 | 0.724 | 101 | 0.898 | 99 | 0.841 | 68 | 0.177 | 61 | 0.105 |

**Supplementary Materials, Table 5. Statistical results of all analysis for Activations count $\alpha_{av}$ over all possible parameters' combinations using a selected set of ROIs.** In bold the significant values ($p < 0.05$)

| *Repeated Correlation across sessions* | | | | |
|---|---|---|---|---|
| ***Parameters' pair*** | ***ΔAVALANCHES' LENGTH $\Delta\lambda_{av}$*** | | ***ΔACTIVATIONS COUNT $\Delta\alpha_{av}$*** | |
| | *r-values* | *p-values* | *r-values* | *p-values* |
| ***$\Theta_{av}$ : $\mu+2\sigma$, $\lambda_{min\_av}$: 5ms*** | ***0.47*** | ***0.0001*** | ***0.33*** | ***0.010*** |
| ***$\Theta_{av}$ : $\mu+2\sigma$, $\lambda_{min\_av}$: 50ms*** | ***0.32*** | ***0.011*** | ***0.26*** | ***0.046*** |
| ***$\Theta_{av}$ : $\mu+2\sigma$, $\lambda_{min\_av}$: 80ms*** | 0.11 | 0.411 | 0.17 | 0.194 |
| ***$\Theta_{av}$ : $\mu+3\sigma$, $\lambda_{min\_av}$: 5ms*** | ***0.44*** | ***0.0004*** | 0.23 | 0.072 |
| ***$\Theta_{av}$ : $\mu+3\sigma$, $\lambda_{min\_av}$: 50ms*** | 0.10 | 0.451 | 0.08 | 0.557 |
| ***$\Theta_{av}$ : $\mu+4\sigma$, $\lambda_{min\_av}$: 5ms*** | ***0.31*** | ***0.014*** | 0.20 | 0.121 |
| ***$\Theta_{av}$ : $\mu+5\sigma$, $\lambda_{min\_av}$: 5ms*** | ***0.25*** | ***0.047*** | 0.17 | 0.181 |

**Supplementary Materials, Table 6. Repeated Correlation results across sessions between both features ($\Delta\lambda_{av}$ , $\Delta\alpha_{av}$) and BCI-scores over all possible parameters' combinations using a selected set of ROIs.** *In bold significant correlations ($p < 0.05$)*

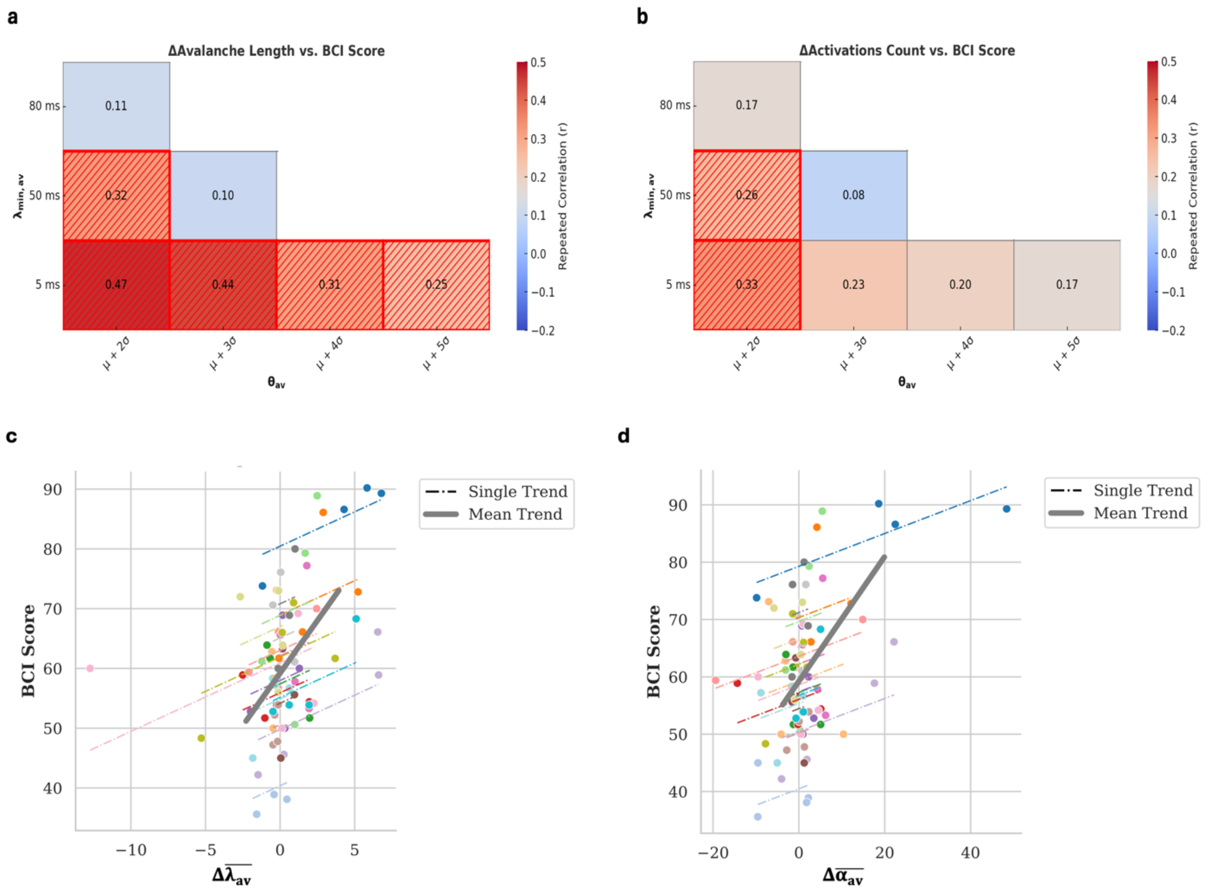

**Supplementary Materials, Figure 3. Repeated correlation and trends across different sessions between BCI score and features changes over a selected set of ROIs**. ***a.*** Mean difference of avalanches' length ($\Delta\lambda_{av}$) and ***b.*** Mean difference activation ($\Delta\alpha_{av}$) across all tested pairs of parameters ($\theta_{av}$, $\lambda_{min\_av}$). Significant correlations ($p<0.05$) are highlighted using different textures. ***c.*** Repeated correlations trend across different sessions between BCI-score and $\Delta\lambda_{av}$ and ***d.*** between $\Delta\alpha_{av}$ and BCI-score. Each coloured dashed line corresponds to one subject while the grey bold line identifies the trend across all the subjects. The pairs of parameters ($\theta_{av}$, $\lambda_{min\_av}$ ) used in ***(c)*** and ***(d)*** were those that achieve the best prediction performance: $\theta_{av}$: $\mu+2\sigma$, $\lambda_{min\_av}$ : 50ms.

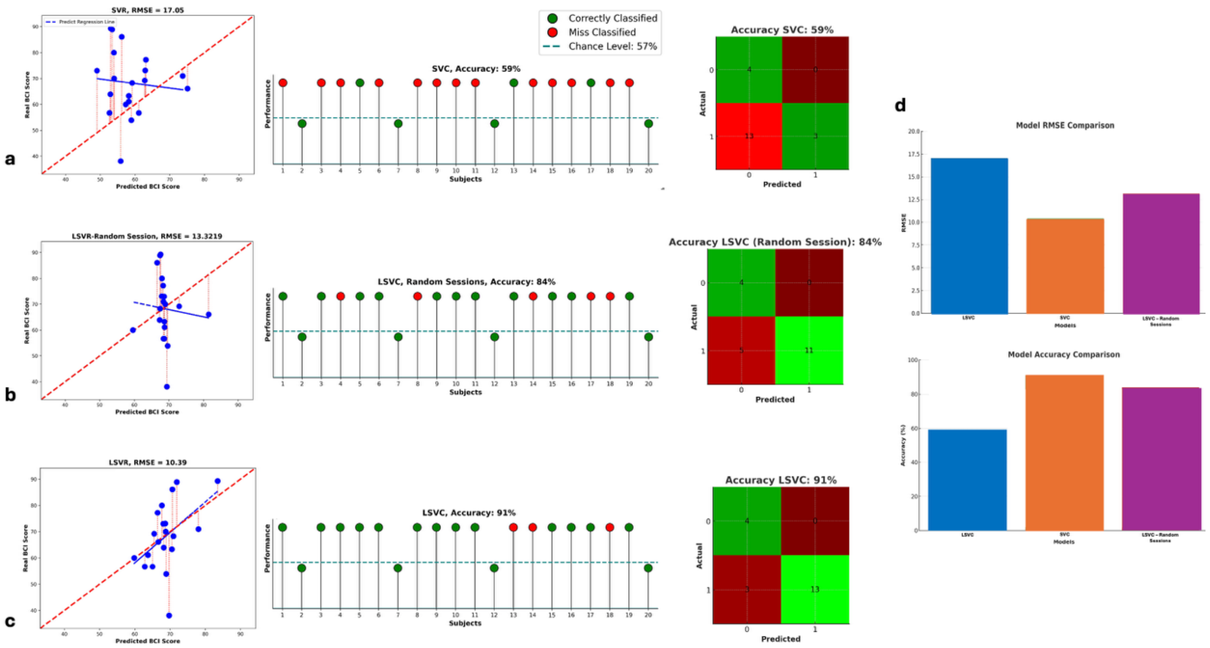

**Supplementary Materials, Figure 4: Predictive Model Results over a selected set of ROIs**.

***a.*** Predictive results using Support Vector Regression (SVR) (left panel) and Support Vector Classification (SVC) (right panel) models.

***b.*** Predictive results using Longitudinal Support Vector Regression (LSVR) (left panel) and Longitudinal Support Vector Classification (LSVC) (right panel) models with sessions in random order.

***c.*** Predictive results using Longitudinal Support Vector Regression (LSVR) (left panel) and Longitudinal Support Vector Classification (LSVC) (right panel) models.

**Left Panel:** Regression Results: Each point represents a subject. Red dashed lines indicate the prediction error between actual and predicted values. The bold red line represents the optimal prediction trend, while the blue lines show individual predicted trends.

**Right Panel:** Classification Results and Confusion Matrix**:** Each ball represents a subject. The dashed horizontal line denotes the classification threshold—subjects above this line are predicted to have control. Green balls indicate correct predictions, while red balls represent misclassifications. In the confusion matrix, green cells show correctly classified subjects, and red cells indicate misclassifications. The intensity of each cell's color corresponds to the number of subjects in that category.

***d.*** Comparison of different models.

**Top:** Comparison of the Root Mean Square Error (RMSE) obtained using different models.

**Bottom:** Comparison of the accuracy performance obtained using different models. SVC (blue), LSVC (orange), and LSVC with random sessions (purple).

All these plots are generated using $\theta_{av}$ $\mu + 2\sigma$ and $\lambda_{min\_av}$ : 50ms, and the best-coupled parameters for prediction.

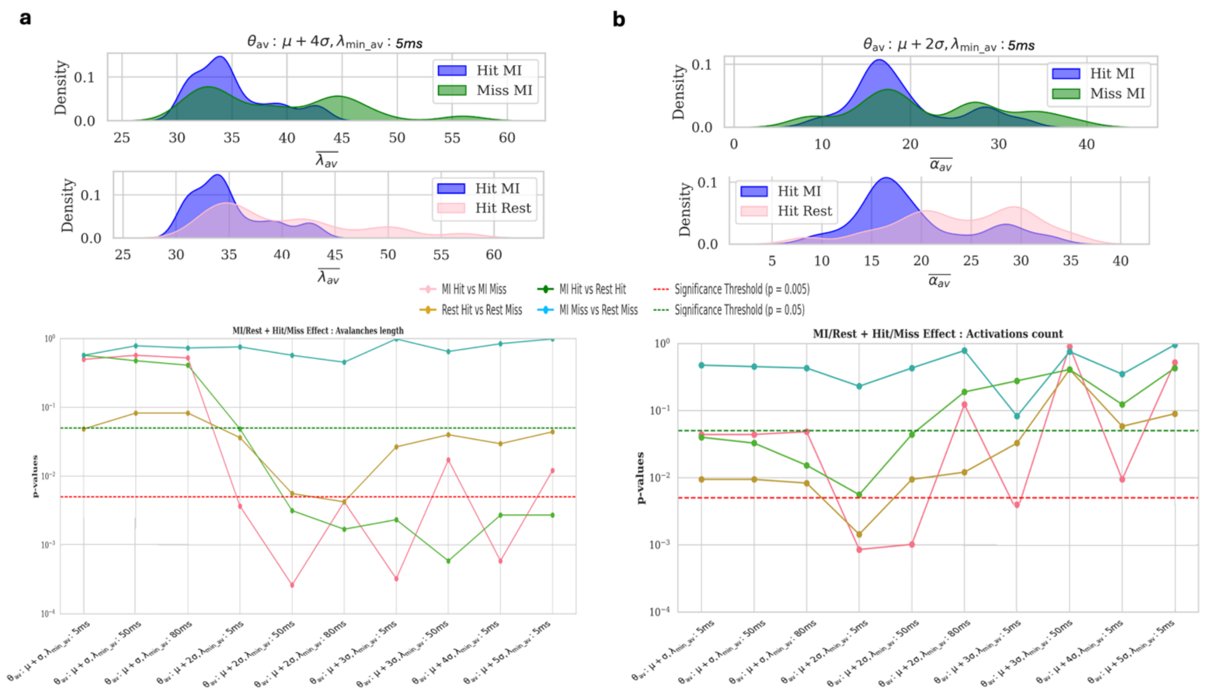

**Supplementary Materials, Figure 5. Analysis of Mean Avalanche Length and Activations in Hit vs. Miss Trials**.

***a.*** Mean Avalanche Length in Hit Trials vs. Miss Trials:

> **Top:** Probability density functions representing: i) Hit Motor Imagery (MI) trials (in blue) vs. Hit Rest trials (in pink) on the left ; ii) Hit MI trials (in blue) vs. Miss MI trials (in green) on the right.
>
> **Bottom:** Trends of p-values from pairwise comparisons of mean avalanche length across brain states (MI vs. Rest) and conditions (Hit vs. Miss) across possible coupled parameters.

Dashed horizontal lines indicate significance thresholds (green: $p < 0.05$: red: $p < 0.001$, Bonferroni-corrected).

***b.*** Activations in Hit Trials vs. Miss Trials:

> **Top:** Probability density functions representing: i) Hit Motor Imagery (MI) trials (in blue) vs. Hit Rest trials (in pink) on the left ; ii) Hit MI trials (in blue) vs. Miss MI trials (in green) on the right.
>
> **Bottom:** Trends of p-values from pairwise comparisons of activation counts across brain states (MI vs. Rest) and conditions (Hit vs. Miss) across possible coupled parameters.

Dashed horizontal lines indicate significance thresholds (green:$p < 0.05$, red: $p < 0.001$, Bonferroni-corrected).

All these analyses are performed only on the last training session.

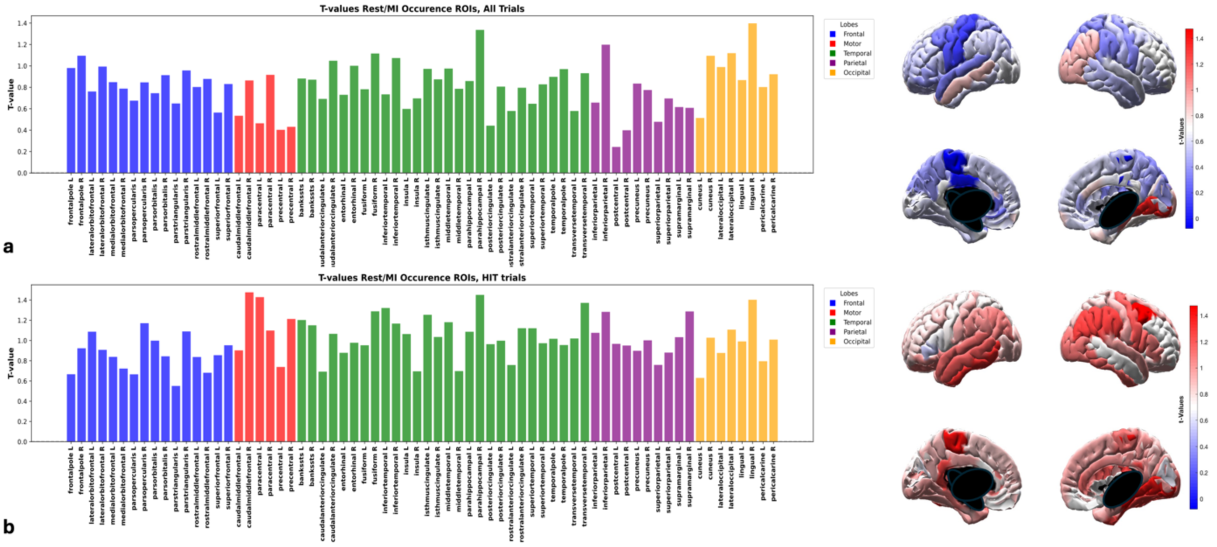

**Supplementary Materials, Figure 6. t-values Comparing Rest and Motor Imagery (MI) Conditions.**
***a.*** t-values calculated for the occurrence of each specific Region of Interest (ROI) across different trials, considering all trials.
***b.*** t-values calculated for the occurrence of each specific ROI across different trials, considering only those trials in which the subject successfully controlled the device.
In the brain plots, the color map reflects the magnitude of the t-values, while in the bar plots, it corresponds to the height of each bar. Brain regions are color-coded as follows: red for the motor cortex, green for the temporal lobe, yellow for the occipital lobe, blue for the frontal lobe, purple for the parietal lobe. All these analyses were performed only on the last training session and one specific parameter combination: $\theta_{av}$: $\mu + 4\sigma$, $\lambda_{min\_av}$: 2.
